# Mitochondrial Lon protease couples substrate translocation to proteolytic activation

**DOI:** 10.64898/2026.06.23.733973

**Authors:** Niko Schenck, Mette Ahrensback Roesgaard, Jan Pieter Abrahams

**Affiliations:** Biozentrum, University of Basel, Basel, Switzerland; Paul Scherrer Institute, Laboratory of Nanoscale Biology, Villigen, Switzerland

## Abstract

Human LonP1 is an ATP-dependent mitochondrial protease that degrades damaged or redundant proteins. Indiscriminate proteolysis by LonP1 is limited through tight coordination of substrate recognition, unfolding, translocation and catalytic cleavage, yet the role of ATP hydrolysis in these individual steps remains unclear. Here, we show that LonP1 binds substrates and cleaves peptide bonds without ATP hydrolysis, whereas degradation of folded proteins strictly depends on ATP-driven unfolding and translocation. Initial substrate binding opens a closed ADP-bound resting state, enabling nucleotide exchange and stimulating ATPase activity. The opening also increases accessibility of the proteolytic chamber, modestly enhancing peptidase activity. Maximal peptidase activity is observed in a transition-state mimic stabilised by ADP·AlF₃, in which substrate is engaged within the translocation channel. Cryo-EM analysis reveals that in this state the proteolytic active sites are no longer occluded, linking ATP-driven substrate translocation to full proteolytic activation. Together, these findings reveal how LonP1 prevents indiscriminate proteolysis during substrate selection by ensuring that efficient proteolysis occurs only in substrate-translocating states.

**Model of the conformational landscape and functional cycle of LonP1:** Schematic overview of LonP1 states and their inter-conversion. State transitions are modulated by substrate, nucleotide occupancy, temperature, and inhibitors. Key distinguishing features include the presence or absence of the lateral gap, nucleotide state, substrate engagement within the A-tunnel, and the handedness of the ATPase (A) domains. Additional indicators include the compactness of the proteolytic (P) domain and the presence of substrate density within the N-terminal (N) domain or at the coiled-coil domain (CCD) as well as the position of a loop within the catalytic centre. The depicted cryo-EM structures represent a model of a continuous conformational landscape and correspond to the closest matching biological states and positions within the reaction cycle, but may also capture transient intermediates or conformations stabilised by experimental conditions. The shown atomic models correspond to the states highlighted in larger font (R-state: PDB 7NGL; P1-state: PDB 7NFY; P2-state: PDB 7NGC; closed LonP1-ADP-substrate: PDB 9CC1).

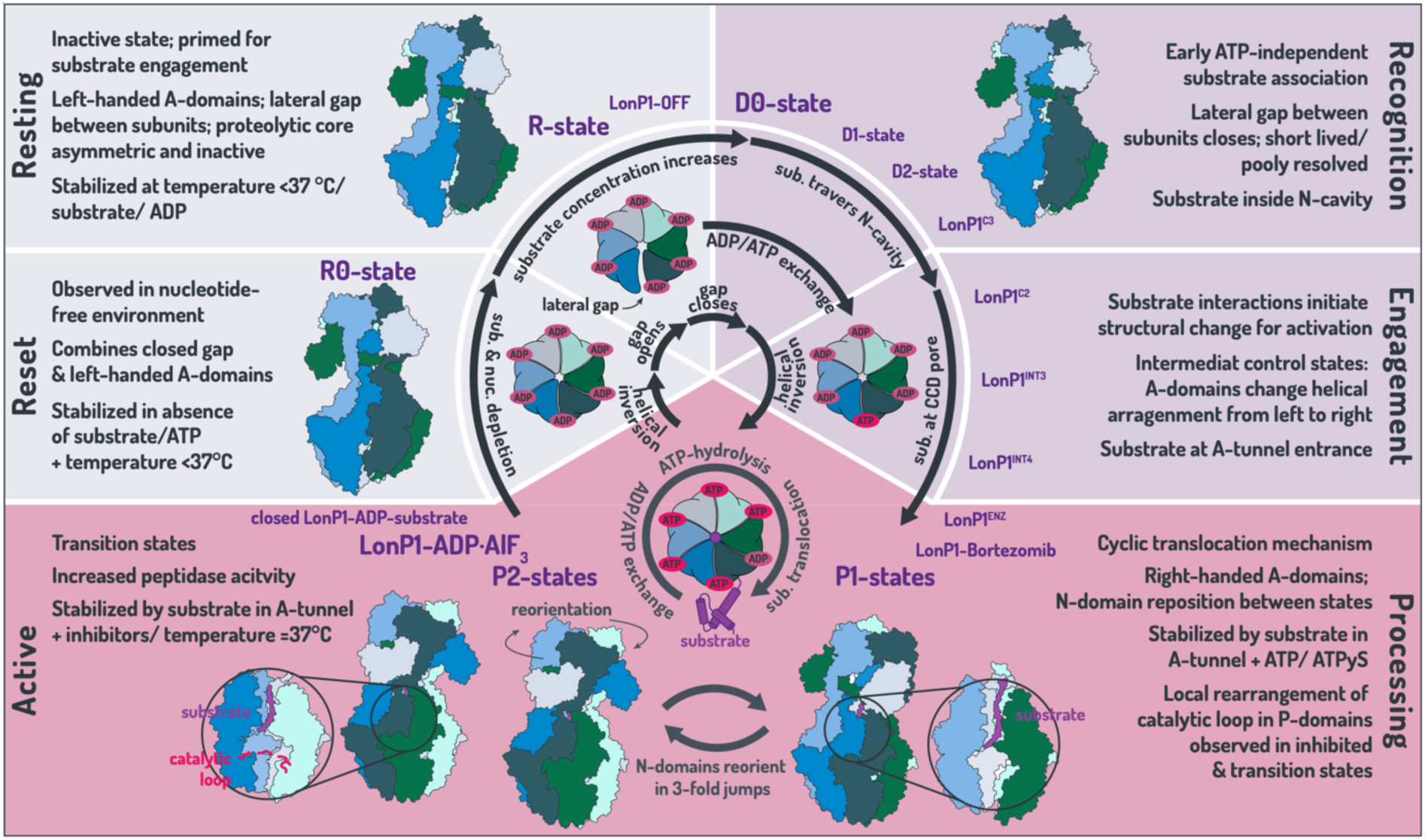

## Introduction

LonP1 is a conserved mitochondrial AAA⁺ protease that maintains proteostasis by unfolding and degrading damaged or regulatory proteins (Pinti et al., 2015; Szczepanowska and Trifunovic, 2022). It couples ATP hydrolysis to substrate unfolding, translocation and proteolytic cleavage within a self-compartmentalised protease chamber (Rotanova et al., 2006; Wlodawer et al., 2022). Recent cryo-EM studies have revealed key features of the LonP1 catalytic cycle. Structural analyses of the human enzyme identified multiple conformational states that support a cyclic substrate-translocation mechanism driven by sequential ATP hydrolysis across the six subunits of the AAA⁺ ring. In these structures, substrate polypeptides are translocated through a central channel formed by the N-terminal and AAA⁺ domains and engaged by conserved pore-loop motifs. These processing states are characterised by a right-handed helical arrangement of the AAA⁺ domains and a closed inter-subunit interface (Mohammed et al., 2022; Shin et al., 2021). Coordinated transitions between ATP-bound, ADP-bound and seam conformations generate a cyclic binding-change mechanism that drives directional substrate translocation, consistent with a Brownian-ratchet-type process. A resting conformation of the enzyme, termed the R-state, is characterised by an ADP-loaded ring in which the A-domains adopt a left-handed helical arrangement, a lateral gap between two subunits, and no visible substrate in the translocation tunnel. The nucleotide-binding site at this interface is poorly ordered, and the density is insufficient to assign ADP or ATP (Mohammed et al., 2022).

While these studies established how ATP hydrolysis powers substrate unfolding and translocation, they primarily address the mechanical aspects of protein processing and leave open how nucleotide state influences substrate recognition and peptide bond cleavage, or how substrate engagement affects ATPase and peptidase activity. Because the proteolytic active sites are sequestered within an internal chamber, productive degradation requires substrate translocation into the protease core. How substrate recognition is coordinated with activation of proteolysis therefore represents a central mechanistic question. In particular, it remains unclear whether proteolytic activity is constitutive or preferentially activated during committed substrate processing.

To address these questions, we combined biochemical and structural approaches to analyse how nucleotide state influences substrate recognition, ATPase activity and proteolysis by LonP1. We compared degradation of a fluorogenic peptide substrate with that of folded protein substrates to distinguish peptide bond cleavage from ATP-driven unfolding and translocation. In parallel, cryo-EM structures of LonP1 in ADP-bound and ADP·AlF₃-stabilised states provide structural snapshots linking nucleotide state, substrate engagement and proteolytic activation.

## Material and Methods

### Lead contact

Jan Pieter Abrahams, Biozentrum, University of Basel, Basel, Switzerland & Paul Scherrer Institute, Laboratory of Nanoscale Biology, Villigen, Switzerland.

### Materials availability

Plasmids for wild-type LonP1 are available upon request. Otherwise, this study did not generate new unique reagents.

### Experimental model and subject details

The human *LONP1* gene sequence encoding the mature protein (residues 125-959, lacking the mitochondrial targeting signal) as well as its proteolytic inactive mutants *LONP1_S855A* and *LONP1_T803V* was used for recombinant expression as previously described (Mohammed et al., 2022). The protein was expressed in *E. coli* BL21 (DE3) cells. Cultures were grown in 2XYT medium at 37 °C until induction. LonP1 purification followed established protocols involving affinity and size-exclusion chromatography (Mohammed et al., 2022).

### ATPase assay

ATPase activity of wild-type LonP1 was measured using a colorimetric ATPase/GTPase assay kit (Sigma-Aldrich, USA, product: MAK113) according to the manufacturer’s instructions. Reactions contained 100 nM LonP1 and 1 mM ATP and were performed in the absence or presence of 100 nM casein. Samples were prepared in ATPase buffer (20 mM HEPES, 100 mM NaCl, 10 mM MgCl2, 0.1 mM EDTA, 1 mM β-mercaptoethanol, 10% glycerol, pH 7.8) and incubated for 15 min at either room temperature or 37 °C. Reactions were terminated by addition of malachite green reagent, and inorganic phosphate release was quantified by measuring absorbance at 620 nm after 30 min using a Tecan Spark® Multimode plate reader. All measurements were performed in triplicate in 96-well plates.

### Microscale Thermophoresis substrate binding assay

Substrate binding to LonP1 was analysed by microscale thermophoresis (MST) to assess nucleotide dependence of substrate recognition. Wild-type LonP1 and the proteolytically inactive variants LonP1_S855A and LonP1_T803V were labelled using RED-tris-NTA 2nd Generation (Monolith His-Tag Labelling Kit, product: MO-L018) according to the manufacturer’s protocol. A 16-step 1:1 serial dilution of casein (starting at 400 µM) was prepared in reaction buffer (50 mM HEPES, 150 mM NaCl, 5 mM MgCl2, pH 7.5), supplemented with either 1% DMSO, or 10 µM CDDO-Me in DMSO, or 10 µM bortezomib in DMSO, and where indicated 2 mM ATP. Labelled protein was pre-incubated with the respective additives for 30 min at room temperature prior to mixing with the casein dilution series, followed by an additional 30 min incubation. MST measurements were performed in triplicate using a Monolith NT.115 instrument (NanoTemper) at 25 °C unless stated otherwise. For temperature-dependent measurements, samples were pre-equilibrated at the indicated temperature (room temperature or 37 °C) prior to mixing and incubation, and measurements were carried out at the corresponding temperature. Data were analysed using MO. AffinityAnalysis for Monolith NT.115 Series Instruments Software. Binding curves were derived from the change in fluorescence between 0.5 and 1.5 s and fitted according to the law of mass action to obtain apparent dissociation constants (KD). Mean values with the corresponding standard error of the mean were calculated from triplicate measurements.

### Peptidase assay

Peptidase activity of LonP1 was assessed using a fluorogenic peptide substrate that can access the proteolytic chamber without requiring ATP-dependent unfolding or translocation (Goldberg et al., 1994). Wild-type LonP1 (185.9 nM) was pre-incubated for 30 min at room temperature in reaction buffer (50 mM HEPES, 150 mM NaCl, 5 mM MgCl2, pH 7.5) in the presence of 12 µM CDDO-Me, 12 µM bortezomib, or an equivalent volume of DMSO. ATP-dependent conditions were established by addition of ATP (1.19 mM), ATP/ADP mixtures, ATPγS, or defined mixtures of ATP and ATPγS at the indicated ratios. Transition-state conditions were generated using aluminium fluoride (AlF₃). For this, LonP1 was pre-incubated with ADP (1 mM) for 20 min, followed by sequential addition of NaF (3.125 mM) and AlCl₃ (0.625 mM) with an additional 20 min incubation. Aluminium fluoride (AlF₃) acts as a phosphate analogue and mimics a transition state of ATP hydrolysis (Braig et al., 2000; Xu et al., 1997). Reactions (42 µL) were transferred to a 96-well microplate (Invitrogen, product: M33089) and initiated by addition of 8 µL of the fluorogenic peptide substrate glutaryl-Ala-Ala-Phe-MNA (final concentration 1.06 mM; Sigma-Aldrich, product: G376). Peptide cleavage was monitored as an increase in fluorescence of 4-methoxy-β-naphthylamide (MNA) over 90 min (excitation 340 nm, emission 420 nm) using a Tecan Spark® Multimode Plate-reader at 25 °C. All measurements were performed in triplicate. Fluorescence signals were background-corrected and normalised. Initial reaction rates were determined from the linear phase of each trace (0-5 min) and are reported as mean with the corresponding standard error of the mean.

### Protease assay

Proteolytic activity of LonP1 towards a protein substrate was assessed using FITC-labelled casein under the same nucleotide and inhibitor conditions as described for the peptidase assay. Wild-type LonP1 was incubated with inhibitors and nucleotides as specified above, and reactions were performed in the same buffer conditions. Reactions were initiated by addition of FITC-casein (final concentration 1.7 µM; Sigma-Aldrich, product: C0528). Where indicated, assays were performed in the presence of 170 µM glutaryl-Ala-Ala-Phe-MNA. Proteolysis was monitored as an increase in fluorescence resulting from fluorescein isothiocyanate (FITC) release over 90 min (excitation 340 nm, emission 420 nm) using a Tecan Spark® Multimode Plate-reader at 25 °C and a 96-well microplate (Invitrogen, product: M33089). All measurements were performed in triplicate. Fluorescence signals were background-corrected and normalised. Initial degradation rates were determined from the linear phase of each trace (0-5 min) and are reported as mean with the corresponding standard error of the mean.

### Protease assay at different temperatures

To control for temperature-dependent effects, casein degradation by LonP1 was additionally assessed by SDS-PAGE instead of using the fluorescence-based assay. Wild-type LonP1 (0.5 µM) was incubated with casein (15 µM) and ATP (3 mM) in reaction buffer (50 mM HEPES, 150 mM NaCl, 5 mM MgCl2, pH 7.5) for 60 min. Samples were collected at the indicated time points and mixed 1:1 with SDS loading buffer (0.13 M Tris-HCl, 0.2 M DTT, 15% glycerol, 0.02% bromophenol blue, 5% SDS, pH 6.8), followed by denaturation at 90 °C for 5 min. Proteins were separated on NuPAGE™ 4-12% Bis-Tris Gel (Thermo Fisher Scientific) using NuPAGE™ MES SDS Running Buffer (Thermo Fisher Scientific, product: NP0002) at 150 V for 60 min. Gels were stained with EZBlue™ Gel Staining Reagent (Sigma-Aldrich), and molecular weights were visualised using Precision Plus Protein Dual Colour Standards (Bio-Rad Laboratories, Inc.).

### Substrate competition assay

To assess the effect of protein substrate binding on peptidase activity, degradation of the fluorogenic peptide substrate glutaryl-Ala-Ala-Phe-MNA was measured in the presence of increasing concentrations of casein. Reactions contained LonP1 (100 nM) and peptide substrate (50 µM), with casein titrated (0, 0.1, 0.3, 1, 3, 10 and 30 µM) in reaction buffer (50 mM HEPES, 150 mM NaCl, 5 mM MgCl2, pH 7.5). ATP (1 mM) was added where indicated. For transition-state conditions, LonP1 was pre-incubated with ADP (2 mM) for 20 min, followed by sequential addition of NaF (3.125 mM) and AlCl₃ (0.625 mM) with an additional 20 min incubation each. Reactions (42 µL) were transferred to a 96-well microplate (Invitrogen, product: M33089) and initiated by addition of LonP1 (8 µL). Peptide cleavage was monitored as an increase in fluorescence of 4-methoxy-β-naphthylamide (MNA) over 90 min (excitation 340 nm, emission 420 nm) using a Tecan Spark® Multimode Plate-reader at 25 °C. For temperature-dependent measurements, reaction components were pre-incubated at 37 °C for 10 min prior to mixing, and assays were performed at 37 °C. All measurements were carried out in triplicate. Fluorescence signals were background-corrected and normalised. Initial reaction rates were determined from the linear phase (4-8 min) and are reported as mean with the corresponding standard error of the mean.

### Structure of LonP1 inhibited by Mg^2+^ ADP and aluminium fluoride

Sample preparation, grid preparation, data acquisition, image processing and model building were described previously (Schenck et al., 2026). Purified LonP1 was subjected to size-exclusion chromatography (Superose 6 Increase 10/300 GL) prior to incubation with ADP (1 mM, 20 min), followed by sequential addition of NaF (3.125 mM) and AlCl₃ (0.625 mM) to generate the ADP·AlF₃ transition-state mimic. Grids were prepared using 3.5 µL sample (1.0 mg/ml) applied to glow-discharged Quantifoil R1.2/1.3 Cu300 grids and plunge-frozen in liquid ethane using a Leica EM GP2 Automatic Plunge Freezer (Leica Microsystems GmbH) (10 °C, 80% humidity, 3.5 s blot time, 80% relative humidity and 75% nitrogen flow). Data were collected on a Thermo Fisher Scientific Titan Krios G4 TEM operated at 300 kV, equipped with a Falcon 4i direct electron detector and a Selectris X TFS Imaging Filter. Movies were acquired using the EPU Software (Thermo Fisher Scientific) in dose-fractionated mode at a pixel size of 0.73 Å. A total electron dose of 40.5 e^−^/Å^2^ was applied over 52 frames (2.7 s exposure). Image processing was performed in cryoSPARC (v4.7.1) (Punjani et al., 2017), including iterative particle picking, 2D classification and heterogeneous refinement. Distinct conformational classes corresponding to P1a- and P2a-like states were identified and refined using non-uniform and local refinement approaches. Model building and refinement were carried out in Coot 0.9.8.96 (Emsley and Cowtan, 2004) and Phenix 1.21 (Adams et al., 2010) using the previously reported P1a structure (PDB: 7NFY) (Mohammed et al., 2022) as a starting model. Nucleotide states were adjusted to ADP·AlF₃, and the model was refined by iterative real-space refinement with standard stereochemical restraints.

### Structure of LonP1 in the absence of additional nucleotides

#### Grid preparation and data acquisition for cryo-electron microscopy

The structure of LonP1 was determined in the absence of added nucleotides, and the impact of substrate protein and mitochondrial DNA on its conformational equilibrium was assessed. The four corresponding datasets (I-IV) were prepared according to Table S1, using a Vitrobot IV (Thermo Fisher Scientific). LonP1 was passed through a size exclusion Superose 6 Increase 10/300 GL column in order to remove aggregates and damaged proteins, prior to incubating in reaction buffer (50 mM HEPES, 150 mM NaCl, 5 mM MgCl2, pH 7.5) for 30 min at room temperature with either LSPas, casein or without any additional components. Cryo-EM datasets were collected on two different transmission electron microscopes under standard low-dose conditions. Datasets I and IV were acquired on a Glacios TEM (Thermo Fisher Scientific) operated at 200 kV and equipped with a Gatan K3 direct electron detector. Movies were recorded at a pixel size of 0.88 Å using SerialEM. Real-time processing and data selection were performed in FOCUS (Biyani et al., 2017). Datasets II and III were collected on a FEI Titan Krios TEM (Thermo Fisher Scientific) operated at 300 kV and equipped with a Gatan K2 Summit detector and a post-column energy filter. Movies were acquired at a nominal pixel size of 0.82 Å using SerialEM and real-time processing and data selection were performed in FOCUS (Biyani et al., 2017).

### Image processing and map generation

Image processing was performed in cryoSPARC v4.1-4.7.1 (Punjani et al., 2017). For dataset I, particles were initially picked from motion-corrected and dose-weighted micrographs using the blob picker. After two rounds of 2D classification, around 17’000 high-quality particles (from 3.3 million initial picks) were used to train a Topaz (Bepler et al., 2019) model for automated particle picking. The resulting 400’000 particles were subjected to iterative 2D classification, yielding approximately 230’000 particles for ab-initio reconstruction (six classes) followed by heterogeneous refinement. Three distinct conformational classes were identified and refined using non-uniform refinement, corresponding to the canonical R-state and two related closed conformations (R1 and R2). Further 3D classification did not reveal additional states. For datasets II-IV, the same Topaz model and processing workflow were applied (after verifying that other picking tools do not yield different results). In the absence of substrate (datasets III and IV), only a single conformational 3D class was observed during heterogeneous refinement and subsequently refined using non-uniform refinement. Local refinement did not result in significant improvements in map resolution or overall quality for any dataset.

### Model building

The atomic model of the new catalytic state was built and refined from dataset IV using Coot 0.9.8.96 (Emsley and Cowtan, 2004) and Phenix 1.21 (Adams et al., 2010). The previously determined R-state of LonP1 (PDB: 7NGL) (Mohammed et al., 2022) served as an initial template. Refinement was performed using an established strategy (Mohammed et al., 2022) consisted of iterative cycles of manual model building in Coot and real-space refinement in Phenix. Refinement included rigid-body fitting of discrete domains followed by global minimisation. Atomic displacement parameters were refined using a grouped B-factor strategy to account for local resolution variations. To maintain structural integrity custom bond restraints were implemented to stabilize the coordination between the P-loop backbone (residues 526-530) and the ADP/ATP ligands. Hydrogen atoms were included during refinement to improve stereochemistry, and side chains were optimised using rotamer correction and automated NQH flips. Model quality was assessed using MolProbity. Structural comparisons were performed using standard superposition procedures in Coot and Phenix.

## Results

### ATP hydrolysis is not required for protein substrate recognition

To determine whether substrate recognition by LonP1 depends on ATP hydrolysis, we measured binding of the model substrate casein using microscale thermophoresis (MST). Fluorophore labelled LonP1 was titrated with increasing concentrations of casein, and binding was quantified from changes in thermophoretic movement (Figure 1). Wild-type LonP1 bound casein in the absence of nucleotides with an apparent affinity in the low micromolar range (KD = 2.95 ± 3.49 µM). Inhibition of the ATPase activity with CDDO-Me produced a similar binding profile (KD = 2.47 ± 1.4 µM), indicating that ATP hydrolysis is not required for substrate association (Figure 1A). Comparable affinities were observed for the proteolytically inactive mutants LonP1_S855A and LonP1_T803V both in the absence and presence of ATP. Likewise, inhibition of the protease active site with bortezomib did not significantly affect casein binding (Figure 1A). Although the MST curves do not define fully saturated binding plateaus and the absolute affinity values should therefore be interpreted with caution, the relative behaviour across all conditions is consistent. Control measurements with unrelated proteins such as lysozyme and eGFP did not reveal detectable interaction with LonP1 (data not shown). Together, these results demonstrate that association of LonP1 with protein substrates occurs independently of ATP hydrolysis and does not require proteolytic activity.

**Figure 1.**
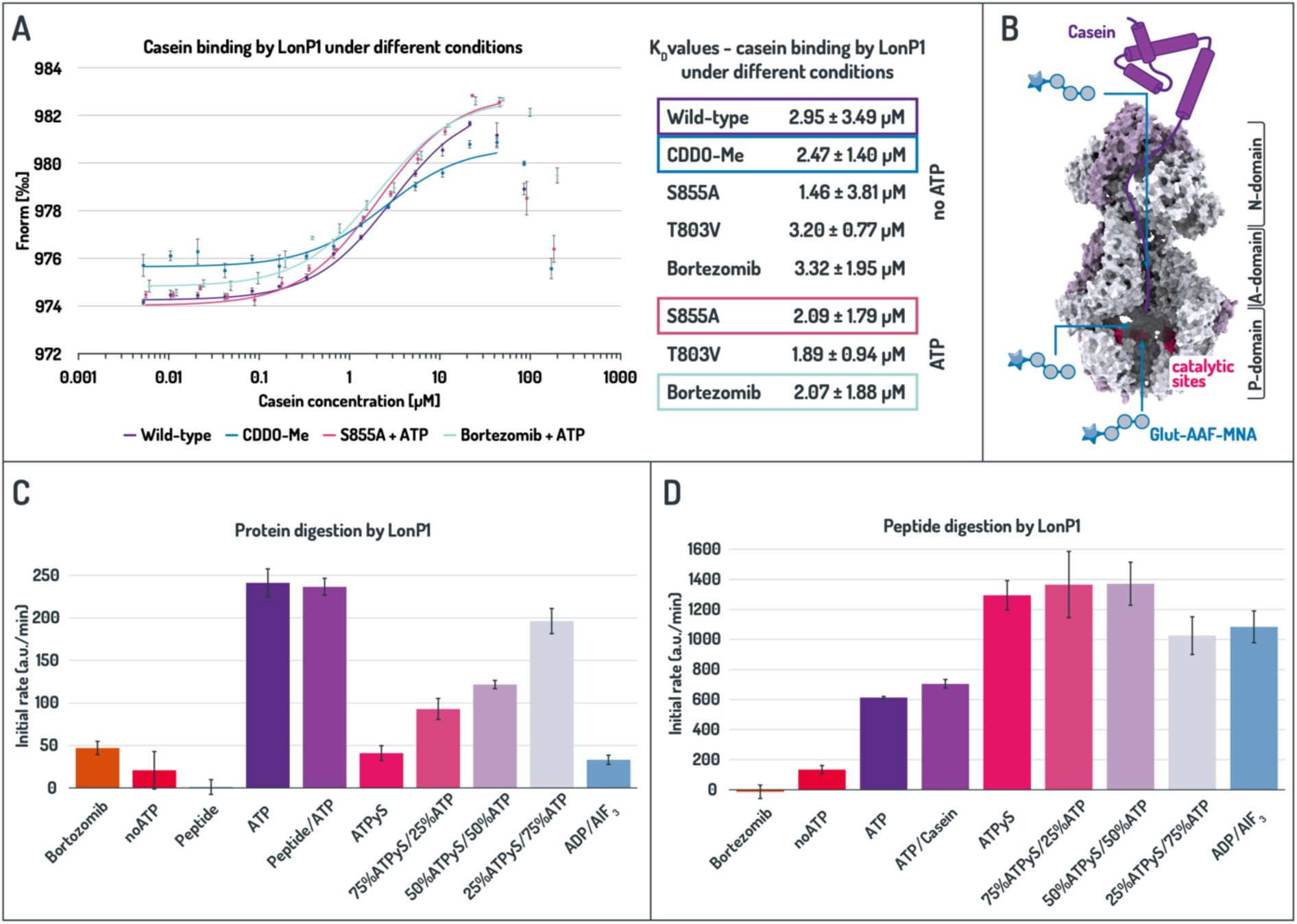
ATP hydrolysis is dispensable for both substrate recognition and peptide bond cleavage by LonP1. LonP1 proteolytic activity and binding dynamics were assessed using the protein substrate casein and the fluorogenic tripeptide Glut-AAF-MNA. **(A)** Representative binding curves of LonP1 variants to casein determined by microscale thermophoresis. Normalized fluorescence (Fnorm) from the initial thermophoretic phase (0.5-1.5 s) was plotted against substrate concentration (mean ± standard error). Apparent dissociation constants (KD) were determined via non-linear regression (outliers at high casein concentrations were excluded). See Figures S2-S6 for individual traces and for all binding curves. **(B)** Schematic representation of substrate access pathways. While protein substrates like casein require N-domain-mediated unfolding and translocation through the A-domain axial pore, the small peptide glutaryl-Ala-Ala-Phe-MNA bypasses the mechanical unfolding machinery to access the proteolytic chamber directly. **(C)** Initial degradation rates of FITC-casein **(D)** and Glut-AAF-MNA under varied nucleotide and inhibitory conditions. Substrate turnover was monitored by fluorescence increase over time; rates were calculated from the linear phase (0-5 min) and are given as mean ± standard error. See Figure S7 for representative fluorescence traces

### Peptide bond cleavage does not require ATP hydrolysis

To determine whether ATP hydrolysis is required for peptide bond cleavage by LonP1, we compared degradation of a small fluorogenic peptide substrate (glutaryl-Ala-Ala-Phe-MNA) with that of the protein substrate FITC-casein under defined nucleotide conditions (Figure 1). The peptide substrate is sufficiently small to diffuse directly into the proteolytic chamber of LonP1, bypassing the requirement for ATP-driven unfolding and translocation, whereas casein must first be actively unfolded and threaded through the central pore.

LonP1 cleaved the peptide substrate in the absence of ATP, as indicated by a gradual increase in fluorescence over time. Although the rate of peptide turnover was reduced compared with ATP-supplemented reactions, the basal activity demonstrates that ATP hydrolysis is not strictly required for peptide bond cleavage. Addition of ATP increased the initial rate of peptide degradation approximately sixfold. Peptidase activity was further enhanced in the presence of the slowly hydrolysable analogue ATPγS and under conditions in which LonP1 was trapped in a transition-state conformation by ADP·AlF₃. In contrast, the protease inhibitor bortezomib abolished peptide cleavage, confirming that the observed fluorescence increase reflects catalytic activity of LonP1 (Figure 1C). In sharp contrast to the peptide substrate, FITC-casein was not degraded in the absence of ATP. Efficient casein degradation required ATP hydrolysis and was abolished when ATP was replaced by ATPγS or ADP·AlF₃, indicating that non-hydrolysable or transition-state-trapping nucleotides cannot support the mechanical steps required for protein substrate processing. Consistently, increasing the fraction of ATP in ATPγS-containing reactions progressively increased the rate of casein degradation (Figure 1D).

Together, these results demonstrate that peptide bond cleavage by LonP1 can occur without ATP hydrolysis when the substrate does not require unfolding or translocation. ATP turnover is therefore essential for the mechanical processes of protein substrate unfolding and translocation but not for the chemical step of peptide bond hydrolysis

### Nucleotide state and substrate binding cooperatively enhance peptidase activity

Recent work proposed that peptidase activity of LonP1 is primarily stimulated by substrate binding rather than by nucleotides (Mindrebo and Lander, 2025). Because our experiments revealed a strong effect of nucleotide state on peptide turnover, we examined how substrate protein and nucleotides jointly influence peptidase activity (Figure 2).

**Figure 2.**
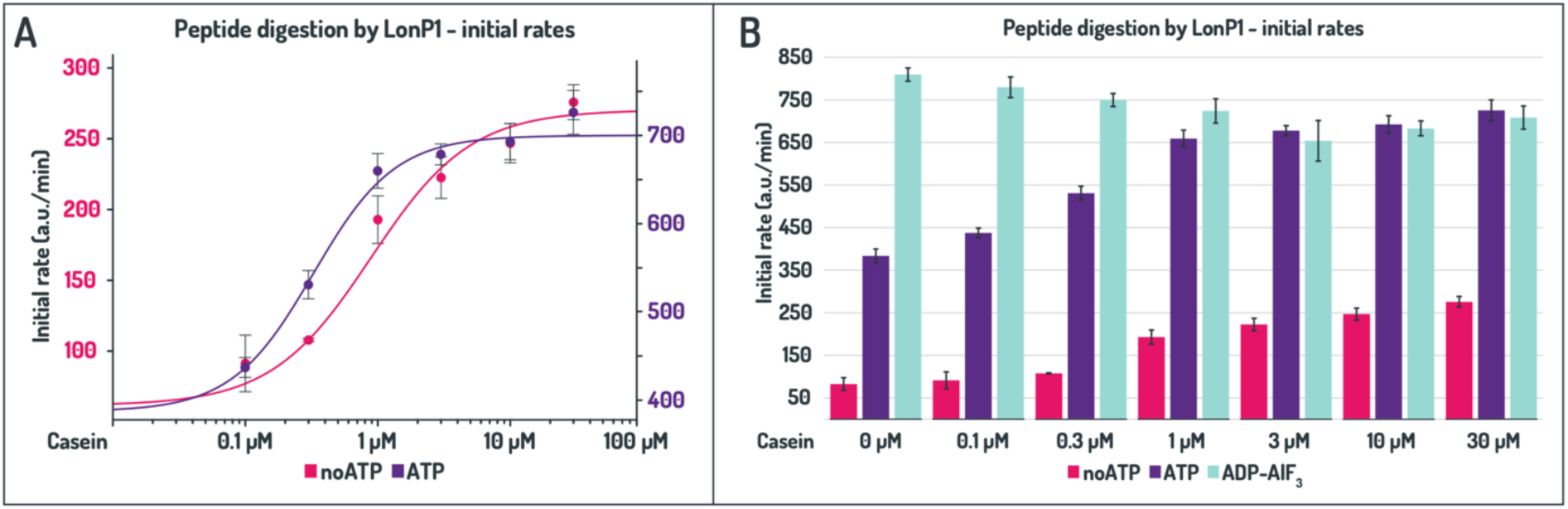
Substrate and nucleotide state cooperatively regulate LonP1 peptidase activity. Initial degradation rates of the fluorogenic peptide substrate glutaryl-Ala-Ala-Phe-MNA by LonP1. Peptidase activity was monitored in the presence of varying concentrations of the protein substrate casein, either in the absence of nucleotides or with the addition of ATP or ADP·AlF₃. **(A)**Allosteric fit of degradation rates. Substrate turnover was determined from the linear phase of fluorescence traces (4-8 min) and plotted on a logarithmic scale (mean ± standard error). **(B)** Comparative analysis of combined initial rates across different nucleotide states.

In the absence of ATP, addition of the substrate protein casein stimulated degradation of the fluorogenic peptide in a sigmoidal manner (Figure 2A). Peptidase activity increased approximately threefold between the lowest and highest casein concentrations tested, indicating co-operative activation by substrate binding. The strongest increase occurred around 1 µM casein, corresponding to substrate saturation at approximately a tenfold molar excess over LonP1 (Figure 2B).

In the presence of ATP, basal peptide turnover was substantially higher. Consistent with substrate-induced activation, ATPase activity of LonP1 increased in the presence of casein (Figure S1), in agreement with previous reports (Mindrebo and Lander, 2025; Mohammed et al., 2022; Shin et al., 2021). Casein still stimulated peptidase activity, but the effect was reduced and saturated at concentrations above 1 µM. In the absence of casein, ATP increased peptide turnover by almost fivefold compared with nucleotide-free reactions, whereas at high casein concentrations the difference between ATP-containing and ATP-free conditions was reduced but remained substantial (Figure 2B).

The strongest activation was observed when LonP1 was trapped in a transition-state conformation by ADP·AlF₃. Under these conditions, peptide turnover was maximal and remained essentially constant across the entire casein titration (Figure 2B), indicating that the enzyme is stabilised in a catalytically activated conformation.

Together, these results show that both substrate binding and nucleotide state modulate the catalytic activity of LonP1, with nucleotide occupancy exerting the dominant effect under our experimental conditions. While substrate binding stimulates peptidase activity, nucleotide-stabilised conformations strongly enhance catalysis and can override additional substrate-dependent activation.

### Structure of an ADP-bound LonP1 resting conformation

To explore the conformational landscape of LonP1, we determined a cryo-EM structure of the enzyme in the absence of added nucleotides or substrate. Despite the absence of exogenous nucleotides, all six nucleotide-binding sites are occupied by endogenous ADP, indicating that LonP1 retains ADP following purification (Figure 3).

**Figure 3:**
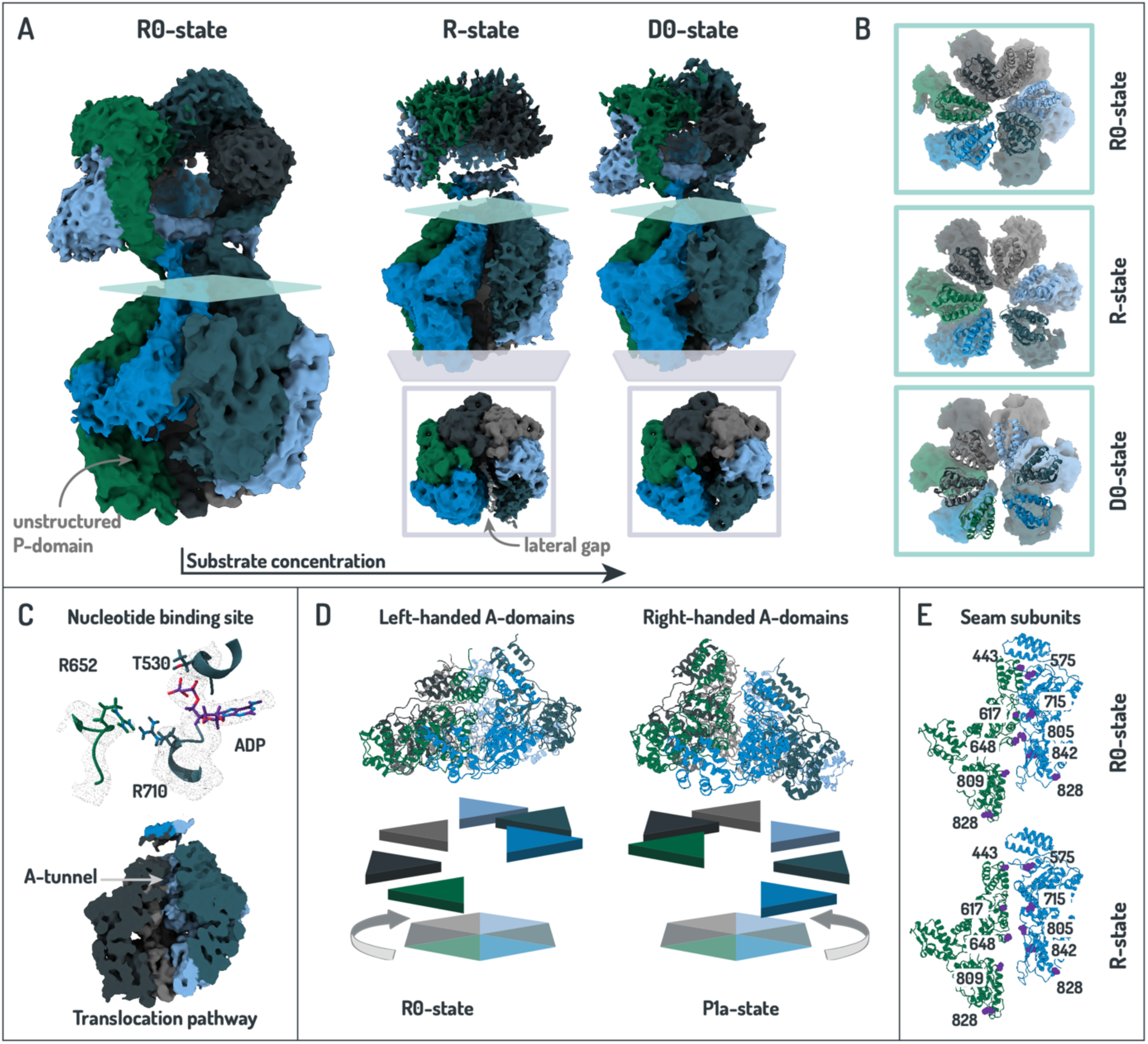
Cryo-EM structure of LonP1 in an ADP-bound resting-like conformation. Cryo-EM analysis of LonP1 in the absence of added nucleotides or substrate reveals a previously uncharacterised conformation. **(A)** Overall architecture of the newly identified conformation and comparison with conformations observed upon substrate addition. Individual subunits of the respective cryo-EM density maps are coloured separately for visual identification. The R0-state is characterised by a partially disordered P-domain adjacent to the seam, whereas the canonical R-state displays a pronounced lateral gap between the A/P-domains together with a diffuse N-terminal assembly. In contrast, the D0-state exhibits a more ordered seam region. **(A)**Comparison of the A-tunnel viewed from the N-terminal cavity toward the proteolytic chamber. **(C)** Nucleotide occupancy and axial channel. Endogenously bound ADP is resolved in all nucleotide-binding pockets (top). No density attributable to a translocating polypeptide is observed within the central axial A-tunnel (bottom). **(D)** Helical organisation of the AAA⁺ domains. The AAA⁺ ring adopts a left-handed pseudo-helical arrangement, distinct from the right-handed organisation observed in processing states. **(E)** Arrangement of the seam subunits at the lateral gap. Neighbouring residues (purple) were used to examine the distance between the two gap subunits more.

This conformation resembles the previously described resting (R) state (Mohammed et al., 2022), with a left-handed pseudo-helical arrangement of the AAA⁺ domains characteristic of ADP-bound LonP1. In contrast to the R-state, which contains a pronounced lateral gap between two neighbouring subunits, the observed conformation displays a closed inter-subunit interface (Figure 3A). This closure results from a rotation of the gap-forming subunits that brings opposing AAA⁺ domains into closer proximity. While the AAA⁺ domains retain a resting-like arrangement and ADP occupancy, the closed interface resembles the more compact architecture observed in processing P-states (Mohammed et al., 2022). Consistent with this compacted architecture, the lateral tunnel appears narrower than in the resting state (Figure 3B, S10). However, in contrast to the P-states, no substrate density is detected within the central A-tunnel (Figure 3C-E). We refer to this closed ADP-bound conformation as the R0-state. In the absence of substrate, LonP1 predominantly adopts this state. Upon addition of the substrate protein casein, the conformational equilibrium shifts toward the canonical, previously reported R-state, characterised by opening of the lateral gap. At higher substrate concentrations, a related intermediate with a narrowed lateral gap becomes more populated (supplemental), which we term the D0 (docking) state. Although both the R0- and D0-states exhibit a closed seam, the D0-state retains a comparatively open A-tunnel and displays more ordered P-domains within the seam subunits (Figure 3A, B).

Together, these observations define a substrate-dependent progression of ADP-bound conformations, in which LonP1 transitions from the closed resting R0-state to the open R-state and, at higher substrate concentrations, further populates the D0-state, priming the enzyme for ATP-driven processing

### Cryo-EM structure of LonP1 trapped in an ATP-hydrolysis transition state

To obtain structural insight into the nucleotide-dependent activation of LonP1, we deter-mined a cryo-EM structure of the enzyme trapped in a transition-state conformation using Mg²⁺, ADP and aluminium fluoride (AlF₃). In this complex, the AlF₃ moiety mimics the γ-phosphate of ATP and stabilises ATPases in a hydrolysis-like configuration (Figure 4C). ADP·AlF₃ was clearly resolved at four inter-subunit catalytic sites, while the remaining two sites contained ADP.

**Figure 4:**
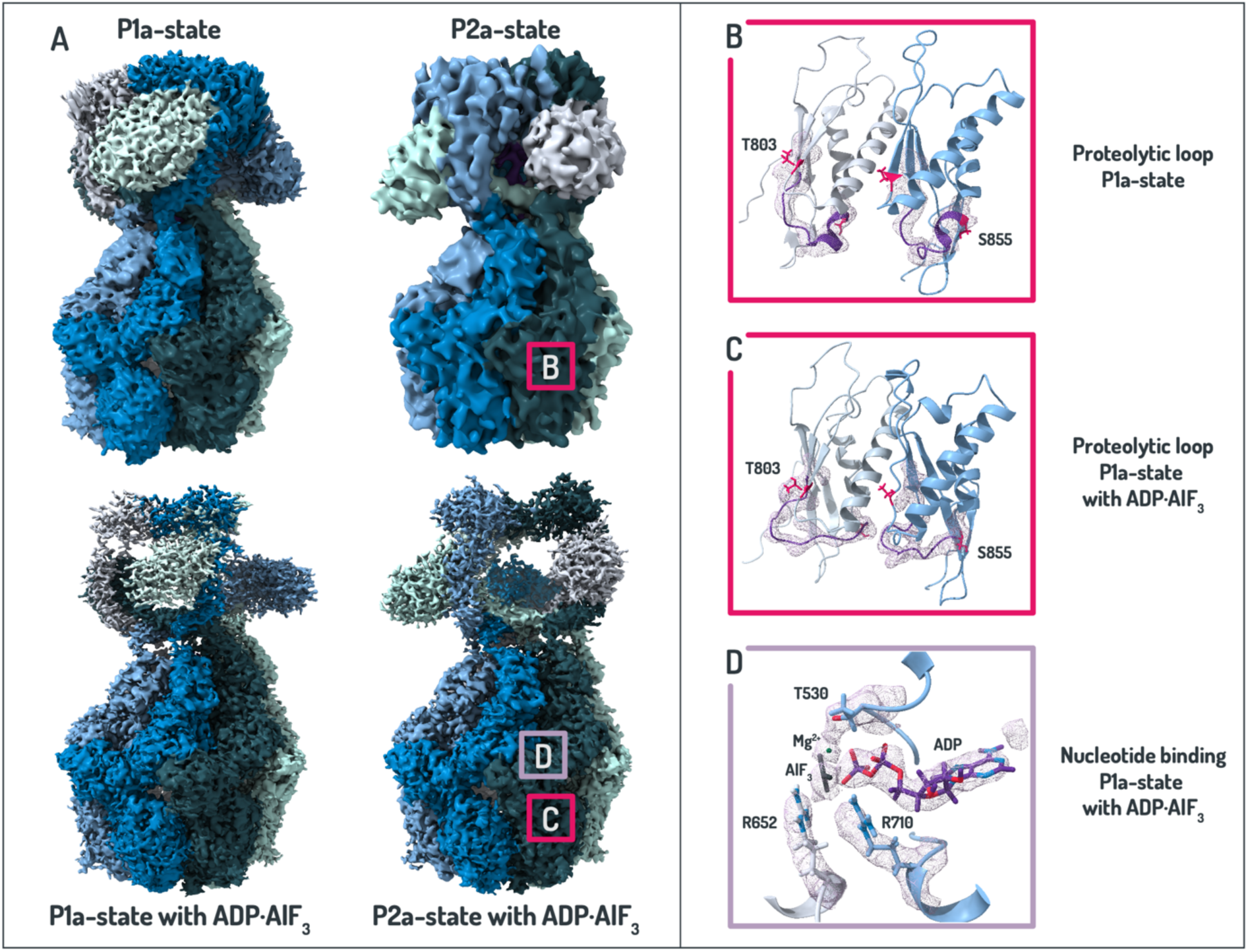
Cryo-EM structure of LonP1 stabilised with ADP·AlF₃ in a transition-like conformation. Cryo-EM analysis of LonP1 in the presence of Mg²⁺, ADP and aluminium fluoride (AlF3) revealed a processing-like conformation with increased structural ordering. **(A)** Overall architecture of the ADP·AlF₃-bound state compared with previously described P1a/P2a processing conformations. The AAA⁺ and protease domains adopt highly similar arrangements, while reduced conformational variability results in improved map resolution. Individual subunits are coloured separately. **(B)** Local rearrangement within the protease domain. A loop spanning residues 844-854 (purple) adopts a distinct conformation relative to P1a/P2a states. Two adjacent subunits are shown, with catalytic residues T803 and S855 indicated in pink. **(C)** Nucleotide-binding pockets. ADP·AlF₃ is re-solved at four catalytic interfaces, while two sites are occupied by ADP, where the resolution does not allow unambiguous assignment of AlF₃.

The structure of LonP1 in the presence of ADP·AlF₃ adopts two conformations closely related to the previously described processing states P1a and P2a (Mohammed et al., 2022) (Figure 4A). In both conformations, the A/P-core is essentially identical, whereas the N-domain assembly differs by an approximately 60° rotation. The AAA⁺ domains form a right-handed helical arrangement with a closed inter-subunit interface, and substrate density is observed within the central A-tunnel. Compared with previously determined processing states, the ADP·AlF₃ complex exhibits a higher degree of structural order within the AAA⁺ and protease rings, resulting in better-resolved overall density and enabling reconstruction to an overall resolution of 2.45 Å, the highest reported for human LonP1.

The improved map quality enabled detailed modelling of the translocating polypeptide within the axial channel. The substrate backbone extends through the VY-pincer region and forms interactions between the substrate carbonyl and the backbone amide of V566, consistent with previously described substrate engagement mechanisms (Mohammed et al., 2022). The density supports modelling of the peptide backbone but does not distinguish a preferred N-to-C or C-to-N orientation, consistent with observations that LonP1 can translocate polypeptides in either direction (Figure 4D) (Schenck et al., 2026).

Despite the overall similarity to previously described processing states, a distinct conformational difference is observed in a loop of the protease domain (residues 844-854) located adjacent to the catalytic residues T803 and S855. This loop adopts a conformation not observed in our earlier structures and is well supported by the density. Similar arrangements have been described as a catalytically active configuration of the protease domain, in which the catalytic serine is no longer occluded by a short 3₁₀ helix (Mindrebo and Lander, 2025; Shin et al., 2021) (Figure 4B). The altered loop conformation therefore provides a structural explanation for the enhanced peptidase activity observed under ADP·AlF₃ conditions.

Together, these observations show that ADP·AlF₃ stabilises LonP1 in a processing conformation with a well-ordered substrate engaged in the central channel and a locally activated protease site.

## Discussion

Our biochemical and structural data support a model in which human mitochondrial LonP1 binds substrates while in a closed ADP-bound resting state. Microscale thermophoresis measurements show that substrate association is independent of ATPase activity, indicating that substrate recognition precedes ATP-driven processing. Similar separation between substrate engagement and ATP-dependent unfolding has been described for other AAA⁺ systems, including the archaeal unfoldase PAN and bacterial Lon homologues (Ibrahim et al., 2017; Smith et al., 2005; Wohlever et al., 2014). Human mitochondrial LonP1 can also bind amyloid and inhibit fibril formation in the absence of ATP (Roesgaard et al., 2026), further supporting the existence of ATP-independent substrate-recognition states. Together, these observations indicate that substrate-recognition surfaces remain accessible within resting conformational ensembles.

Cryo-EM analysis indicates that initial substrate binding promotes opening of the closed ADP-bound R0 resting state. This conformational transition likely facilitates nucleotide exchange through opening of the A-domain interface, thereby enabling ADP/ATP exchange and subsequent ATPase activity. At the same time, opening of the lateral translocation channel and parting of the corresponding P-domains, increases accessibility of the proteolytic chamber, which may explain the modestly enhanced peptidase activity prior to substrate engagement within the translocation pathway. To further investigate the structural basis for the reduced peptidase activity of the barely active R0-state relative to the more proteolytic open R- and closed D0 conformations, we compared the interfaces between adjacent P domains containing the proteolytic sites. Whereas most interfaces were highly similar across the different states, with their catalytic S855 residues buried inside the proteolytic chamber, the seam-adjacent subunit exposed S855 to the exterior. In the R0-state, however, this domain was markedly less ordered, which may contribute to the reduced catalytic competence of the R0 conformation.

ATP hydrolysis subsequently drives substrate unfolding and translocation, whereas peptide bond cleavage itself does not require ATP turnover. Small fluorogenic peptides are therefore cleaved in the absence of ATP, consistent with their ability to access the proteolytic chamber without mechanical processing. However, peptidase activity reaches its maximum in transition-state-like conformations in which substrate is engaged within the translocation channel and the proteolytic sites are no longer occluded. Given the sequential nature of ATP hydrolysis around the AAA⁺ ring, these observations support a model in which ATP-driven substrate translocation is coupled to transient sequential activation of proteolytic sites within the proteolytic chamber (Mohammed et al., 2022; Shin et al., 2021).

Recent work has proposed that increasing temperature favours closed, catalytically competent conformations and that substrate binding is a primary activator of proteolytic activity in LonP1 (Mindrebo and Lander, 2025). Consistent with this, we observe that elevated temperature increases ATPase, protease and peptidase activity while reducing substrate binding, and that substrate binding enhances peptidase activity (Figure S1).

Our data indicate that nucleotide state provides an additional layer of regulation of proteolysis, with activity further increased under ATP-bound conditions and peaking in a transition-state mimic. This behaviour is consistent with observations in bacterial Lon proteases, where ATP binding enhances peptidase activity (Goldberg et al., 1994; Thomas-Wohlever and Lee, 2002). Trapping LonP1 with ADP·AlF₃ stabilises processing-like conformations with a right-handed helical arrangement, substrate density in the axial channel and a highly ordered protease domain. A loop adjacent to the catalytic residues adopts a conformation associated with proteolytic activation (Mindrebo and Lander, 2025; Shin et al., 2021), providing a structural explanation for the enhanced peptidase activity observed under transition-state conditions. Importantly, the ADP·AlF₃-stabilised state likely exaggerates the simultaneous occupancy of transition-state conformations relative to the native catalytic cycle. Whereas four of the six nucleotide-binding sites adopt transition-state-like configurations in our structure, sequential ATP hydrolysis during active turnover is expected to involve only a subset of interfaces at any given time (Mohammed et al., 2022; Shin et al., 2021). The observed structure therefore most likely represents a stabilised processing ensemble enriched in catalytically competent states rather than a single instantaneous configuration sampled during substrate translocation.

We also observed enhanced peptidase activity in the presence of ATPγS, likely reflecting prolonged occupancy of hydrolysis-associated conformations due to the reduced rate of γ-phosphate cleavage. Unlike ADP·AlF₃, which stabilises a transition-state mimic, ATPγS may increase the lifetime of transition-state-adjacent conformations, thereby increasing the probability of sampling catalytically competent states (Mohammed et al., 2022).

Taken together, our results indicate that ATP hydrolysis in LonP1 primarily powers substrate unfolding and translocation while restricting maximal proteolytic activation to substrate-translocating states. This organisation permits ATP-independent substrate surveillance while ensuring that efficient proteolysis occurs only after substrate engagement within the translocation pathway. Rather than functioning as a simple on-off switch, nucleotide binding and hydrolysis regulate transitions between conformational states that differ in catalytic competence. In this view, the functional output of LonP1 emerges from a dynamic conformational landscape in which substrate binding, nucleotide state and translocation cooperatively control protease activation. Future work will be required to determine how substrate occupancy, nucleotide turnover and mitochondrial stress conditions redistribute these conformational ensembles during active proteolysis in vivo.

## Acknowledgements

We thank Mohamed Chami, Carlos Fernandez Rodriguez and Rémi Ruedas for supporting cryo-EM data collection at the Biozentrum BIO-EM lab; Inayathulla Mohammed for advice on data processing; Timothy Sharpe and Alexander Schmidt for the biophysics and MS facility at the Biozentrum Basel.

## Supplemental data

Recent work by Mindrebo and Lander showed that temperature strongly influences the conformational landscape of human LonP1, yielding structural states that differ from those observed under room-temperature conditions (Mindrebo and Lander, 2025). Because most of our experiments were performed at room temperature, we examined how elevated temperature affects LonP1 activity and substrate interaction to assess whether temperature differences could explain discrepancies between the two studies.

Biochemical measurements revealed a clear stimulation of LonP1 activity at higher temperature. The ATPase rate increased from 33.8 ± 0.9 µM ATP hydrolysed per minute at room temperature to about 55.5 ± 5.4 µM per minute at 37 °C. Independent of temperature, the presence of the substrate protein casein increased the ATPase activity of LonP1 by approximately 10 µM ATP per minute (Figure S1B).

In addition, proteolytic turnover of the protein substrate casein was accelerated with digestion that required approximately 14 minutes at room temperature was completed in 4 minutes at 37 °C (Figure S1C).

Temperature also altered the relative contributions of nucleotides and substrate to peptide turnover (Figure S1D). At room temperature, ATP strongly stimulated peptide cleavage, increasing activity by up to almost 5-fold in the absence of casein. In contrast, at 37 °C ATP had little additional effect when casein was absent or present below 1 µM, as basal activity was already elevated. Only at higher casein concentrations did ATP modestly increase turnover. Substrate addition continued to stimulate peptide cleavage in a sigmoidal manner, but the magnitude of stimulation was substantially reduced compared with room temperature. At 37 °C, substrate increased activity by approximately 1.4-fold in the absence of ATP and 1.6-fold in its presence, whereas the corresponding increases at room temperature were approximately 3-fold and 2-fold, respectively. Despite these changes, the transition-state mimic ADP·AlF₃ remained inhibitory for protein degradation (data not shown) but produced the highest peptidase activity and remained insensitive to casein titration.

Interestingly, substrate binding affinity decreased at elevated temperature. While binding at room temperature occurred with an apparent KD of approximately 1.7 µM, the affinity decreased to about 4.4 µM at 37 °C (Figure S1A). If LonP1 and substrate were premixed at 37 °C prior to incubation (instead of first bringing both separately to the target temperature and then incubating them together at 37 °C), the apparent dissociation constant increased further (data not shown). Thus, increased temperature enhances catalytic activity while weakening substrate binding.

Together, these observations suggest that physiological temperature shifts the conformational equilibrium of LonP1 toward nucleotide-activated, catalytically competent states that favour enzymatic turnover but are less permissive for initial substrate binding. Increased ATPase and protease activity further indicate that conformational transitions occur more rapidly under these conditions.

### Structure of LonP1 in the absence of additional nucleotides

To systematically assess the determinants of LonP1 conformational states, cryo-EM datasets were collected under varying conditions, including the presence or absence of substrate (casein), mitochondrial single-stranded DNA (LSPas), and exogenous nucleotides. Comparative analysis across these conditions revealed a distinct closed R0 conformation that was consistently enriched in the absence of added nucleotides. In contrast, particles adopting the previously described R-state or a related closed D1 conformation were also observed in the presence of substrate. These findings indicate that the nucleotide state is a primary determinant of the conformational distribution of LonP1, with nucleotide-free conditions favouring the closed R0 state. Neither the addition of LSPas alone nor in combination with substrate measurably altered the conformational landscape of LonP1. Consistently, no additional cryo-EM density could be assigned to bound DNA. While previous biochemical studies reported an interaction between LonP1 and LSPas that is enhanced by substrate (Liu et al., 2004), such interactions were not resolved in our structural analysis. This may reflect dissociation during grid preparation or the presence of weak, transient, or heterogeneous DNA binding that is averaged out during reconstruction.

---

**Figure S1:**
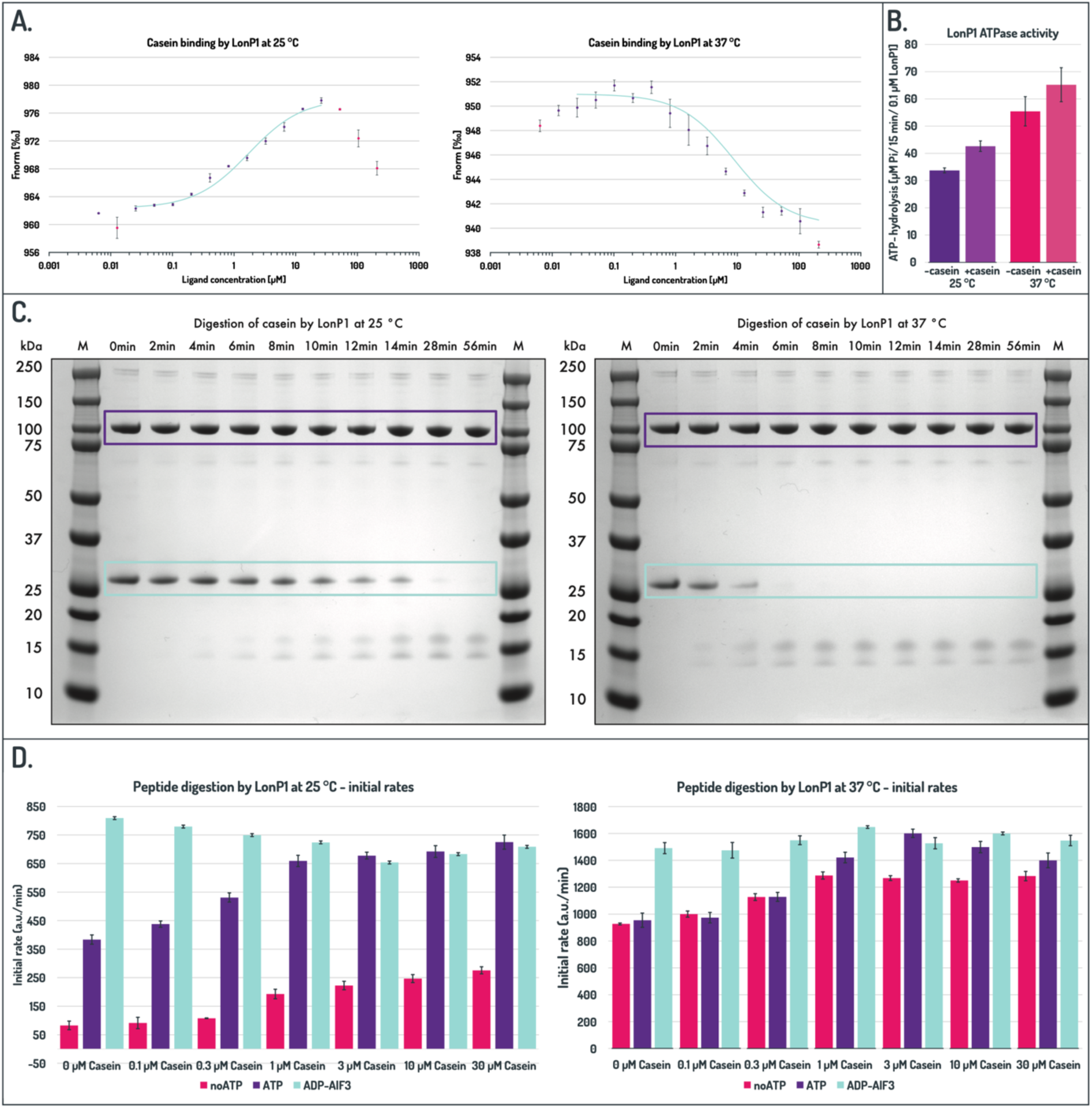
Temperature-dependent effects on LonP1 activity and substrate interaction. The ATPase, protease and peptidase activity as well as the ability to bind a substrate protein was analysed at 37 °C and room temperature (25 °C). **A.** Substrate binding measured by microscale thermophoresis. Binding curves for fluorophore-labelled LonP1 titrated with increasing concentrations of casein are shown (mean ± standard error). Normalized fluorescence (Fnorm) from the initial thermophoretic phase (0.5-1.5 s) was plotted against substrate concentration (mean ± standard error). Apparent dissociation constants (KD) were determined via non-linear regression (outliers at high casein concentrations are indicated in pink). Left, casein binding at 25 °C (KD = 1.72 ± 5.56 µM) and to the right casein binding at 37 °C (KD = 4.44 ± 4.76 µM). **B.** ATPase activity quantified by malachite green assay (Sigma-Aldrich, USA, product: MAK113). Released inorganic phosphate after 15 min incubation was determined by a colorimetric reaction according to the manufacturer’s instructions (mean ± standard error). **C.** Proteolytic turnover of casein analysed by SDS-PAGE at indicated time points. Bands corresponding to casein and LonP1 are marked in cyan and purple, respectively. **D.** Peptidase activity was analysed using the fluorogenic peptide glutaryl-Ala-Ala-Phe-MNA. The effect of increasing concentrations of casein was assessed in the absence and presence of ATP or ADP·AlF₃ and at different temperatures. Substrate turnover was monitored as an increase in fluorescence over time, the initial degradation rates are presented as mean ± standard error.

**Figure S2:**
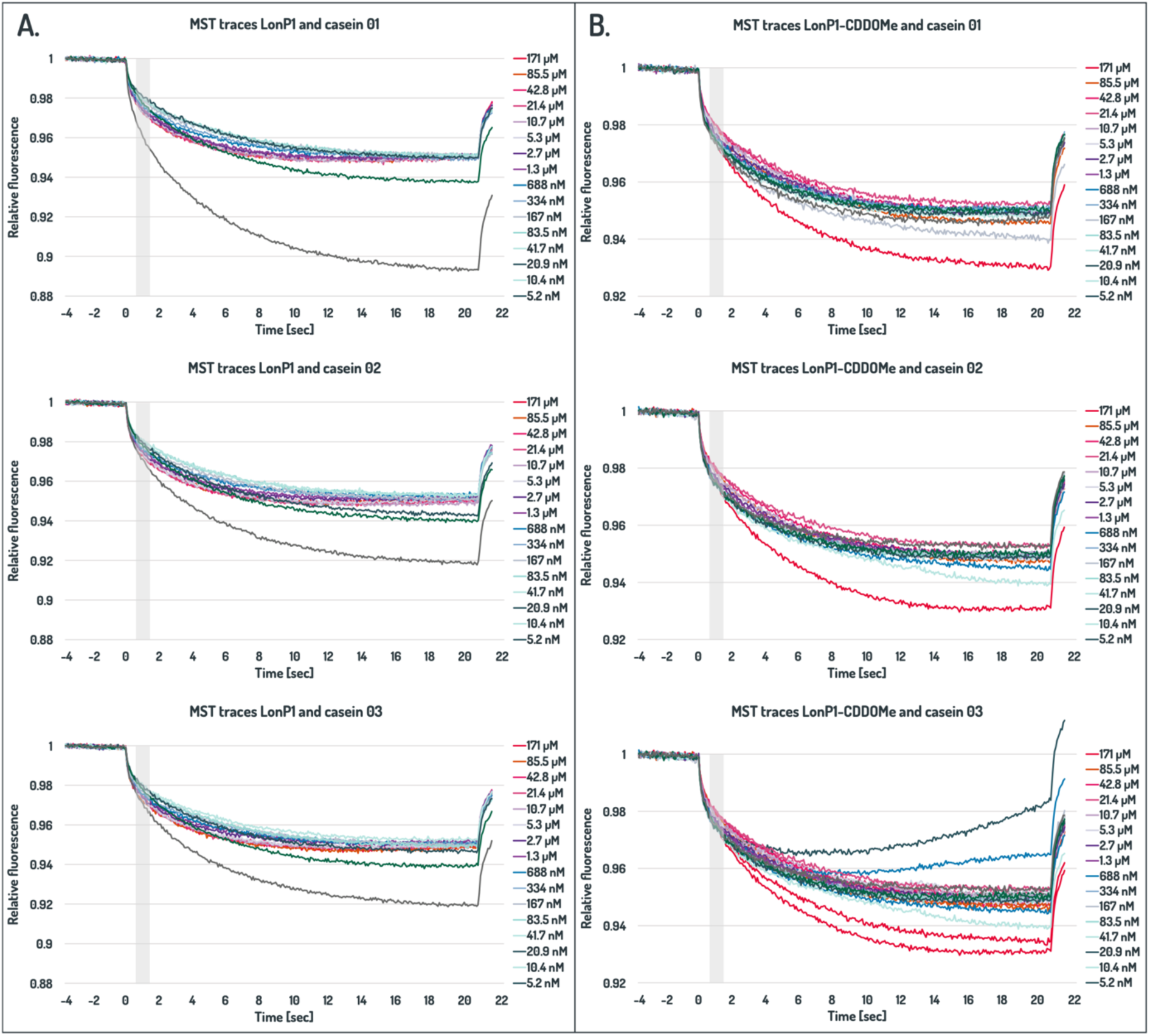
Substrate binding by wild-type LonP1 in the presence and absence of an ATPase inhibitor. Initial substrate recognition by LonP1 was analysed by microscale thermophoresis (MST). N-terminally fluorophore-labelled LonP1 was incubated with increasing concentrations of the substrate casein in the absence or presence of the ATPase inhibitor CDDO-Me (or corresponding buffer control). Thermophoresis was monitored as changes in fluorescence before laser heating, during IR-laser application, and after heating. Relative fluorescence changes were calculated from the initial thermophoretic phase (0.5-1.5 s; grey area). Measurements were performed in triplicate. The MST traces were used to calculate the apparent binding affinities (Figure S6) **A.** Binding of casein to labelled wild-type LonP1 without additional components. **B.** Effect of CDDO-Me on casein recognition by LonP1

**Figure S3:**
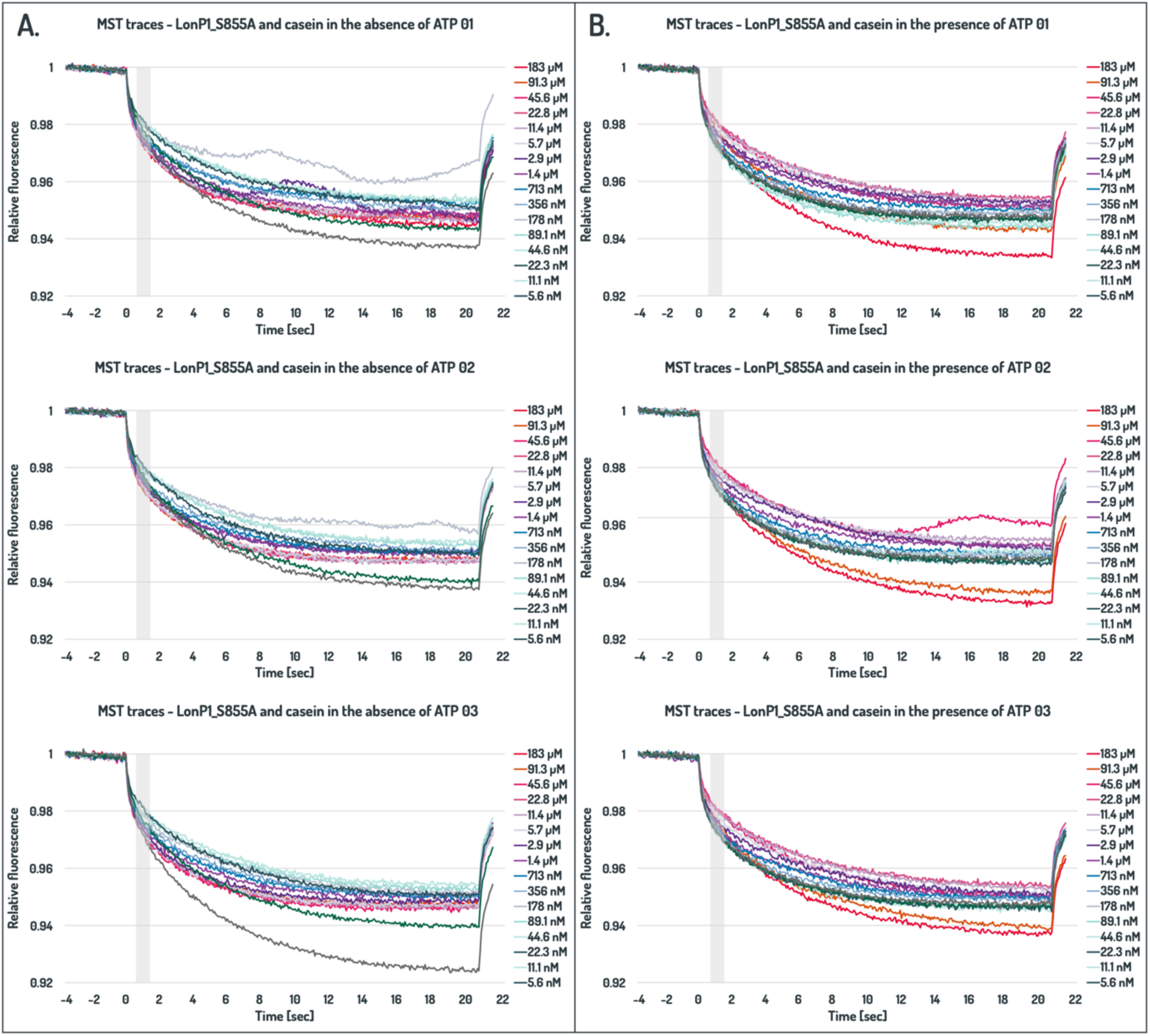
Substrate binding by the proteolytically inactive mutant LonP1_S855A in the presence and absence of ATP. Initial substrate recognition by LonP1_S855A was analysed by microscale thermophoresis (MST). N-terminally fluorophore-labelled LonP1_S855A was incubated with increasing concentrations of the substrate casein in the absence or presence of ATP. Thermophoresis was monitored as changes in fluorescence before laser heating, during IR-laser application, and after heating. Relative fluorescence changes were calculated from the initial thermophoretic phase (0.5-1.5 s; grey area). Measurements were performed in triplicate. The MST traces were used to calculate the apparent binding affinities (Figure S6). **A.** Binding of casein to labelled LonP1_S855A in the absence of ATP. **B.** Effect of ATP on casein recognition by LonP1_S855A.

**Figure S4:**
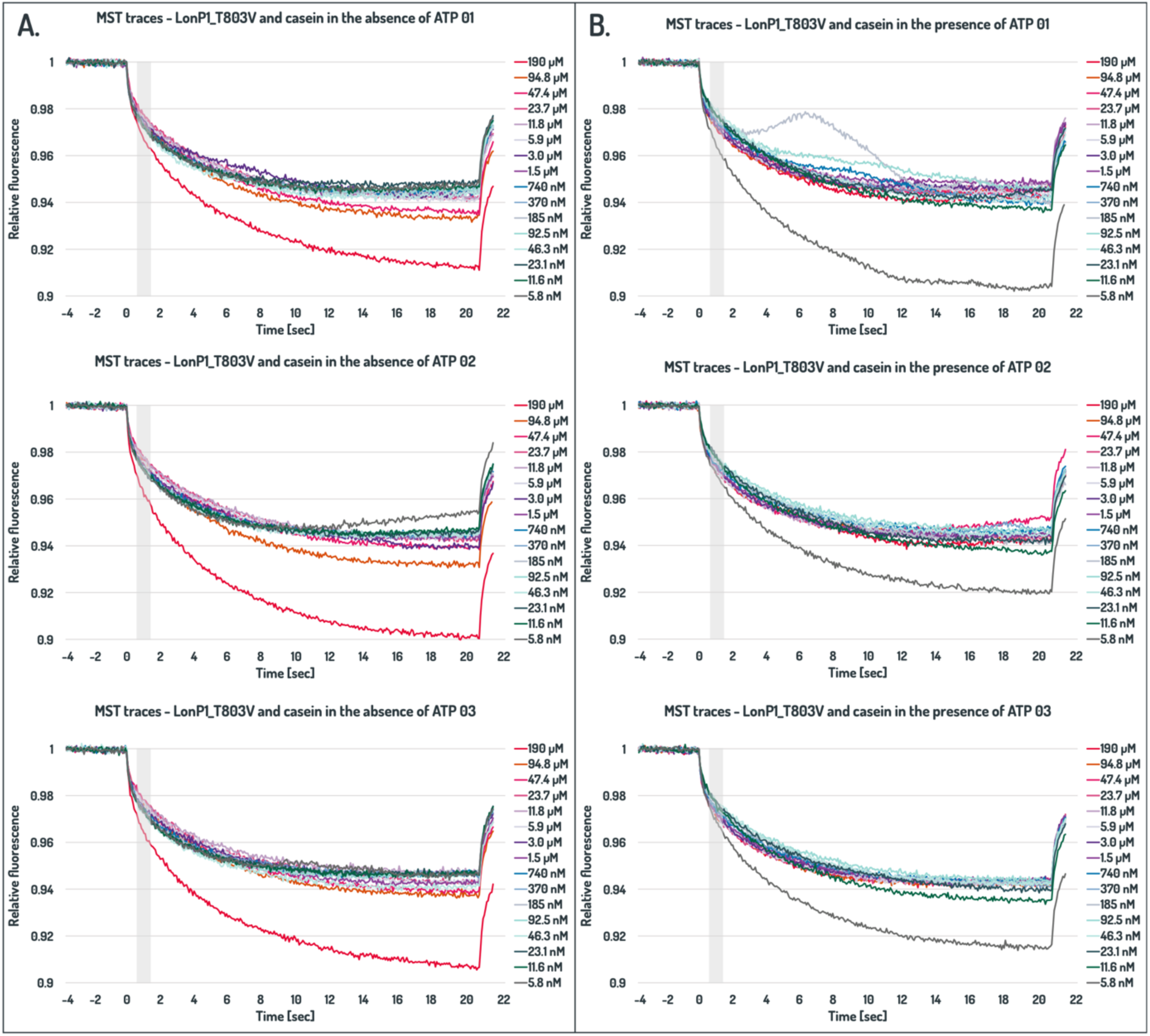
Substrate binding by the proteolytically inactive mutant LonP1_T803V in the presence and absence of ATP. Initial substrate recognition by LonP1_T803V was analysed by microscale thermophoresis (MST). N-terminally fluorophore-labelled LonP1_T803V was incubated with increasing concentrations of the substrate casein in the absence or presence of ATP. Thermophoresis was monitored as changes in fluorescence before laser heating, during IR-laser application, and after heating. Relative fluorescence changes were calculated from the initial thermophoretic phase (0.5-1.5 s; grey area). Measurements were performed in triplicate. The MST traces were used to calculate the apparent binding affinities (Figure S6). **A.** Binding of casein to labelled LonP1_T803V in the absence of ATP. **B.** Effect of ATP on casein recognition by LonP1_T803V.

**Figure S5:**
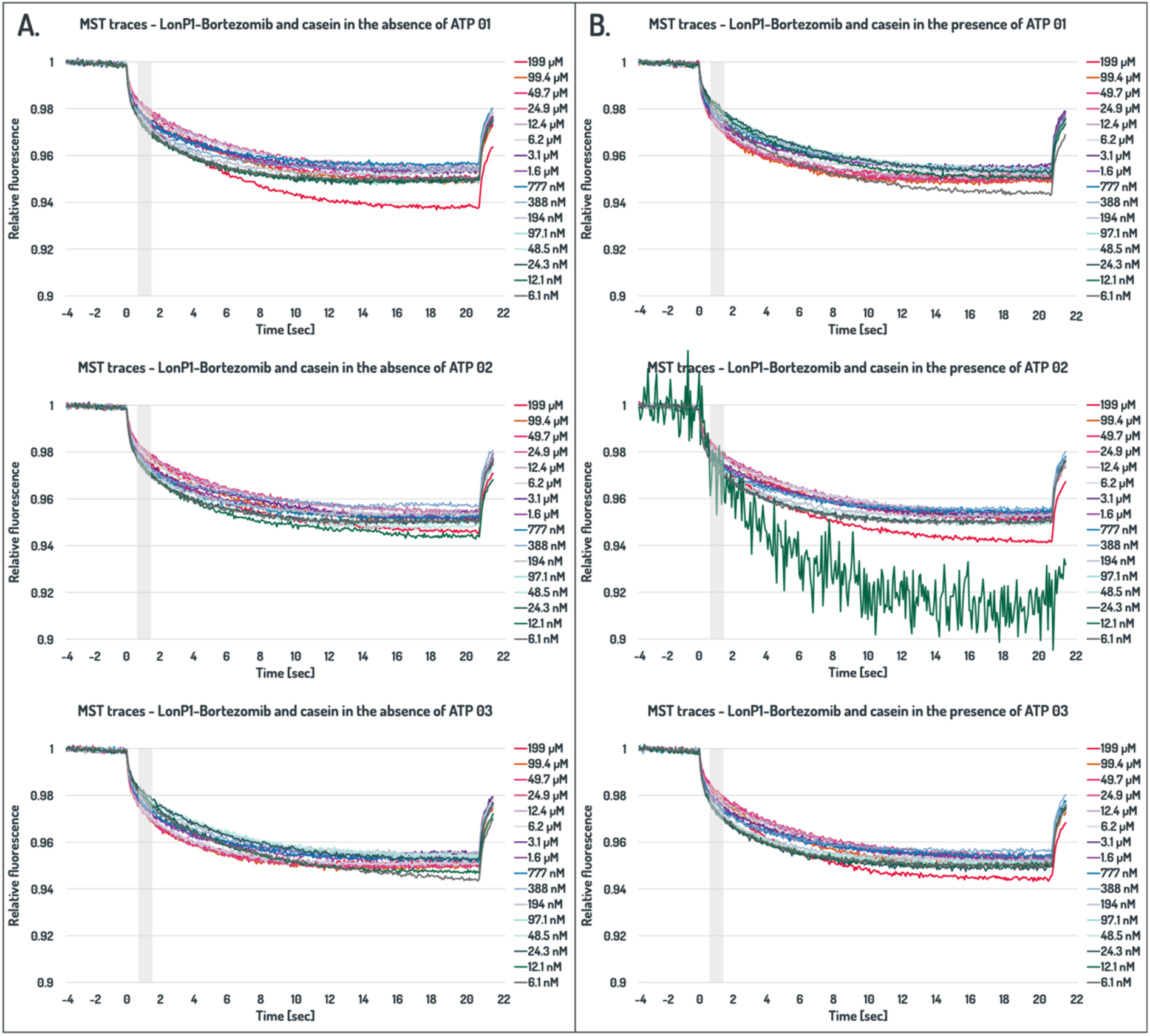
Substrate binding by proteolytically inhibited LonP1 in the presence and absence of ATP. Initial substrate recognition by wild-type LonP1 was analysed by microscale thermophoresis (MST). N-terminally fluorophore-labelled LonP1 was pre-incubated with the active-site inhibitor bortezomib and subsequently incubated with increasing concentrations of the substrate casein in the absence or presence of ATP. Thermophoresis was monitored as changes in fluorescence before laser heating, during IR-laser application, and after heating. Relative fluorescence changes were calculated from the initial thermophoretic phase (0.5-1.5 s; grey area). Measurements were performed in triplicate. The MST traces were used to calculate the apparent binding affinities (Figure S6). **A.** Binding of casein to bortezomib-inhibited LonP1 in the absence of ATP. **B.** Effect of ATP on casein recognition by bortezomib-inhibited LonP1. The MST trace at 12.1 nM ligand (green) in measurement 02 was excluded from Fnorm determination.

**Figure S6:**
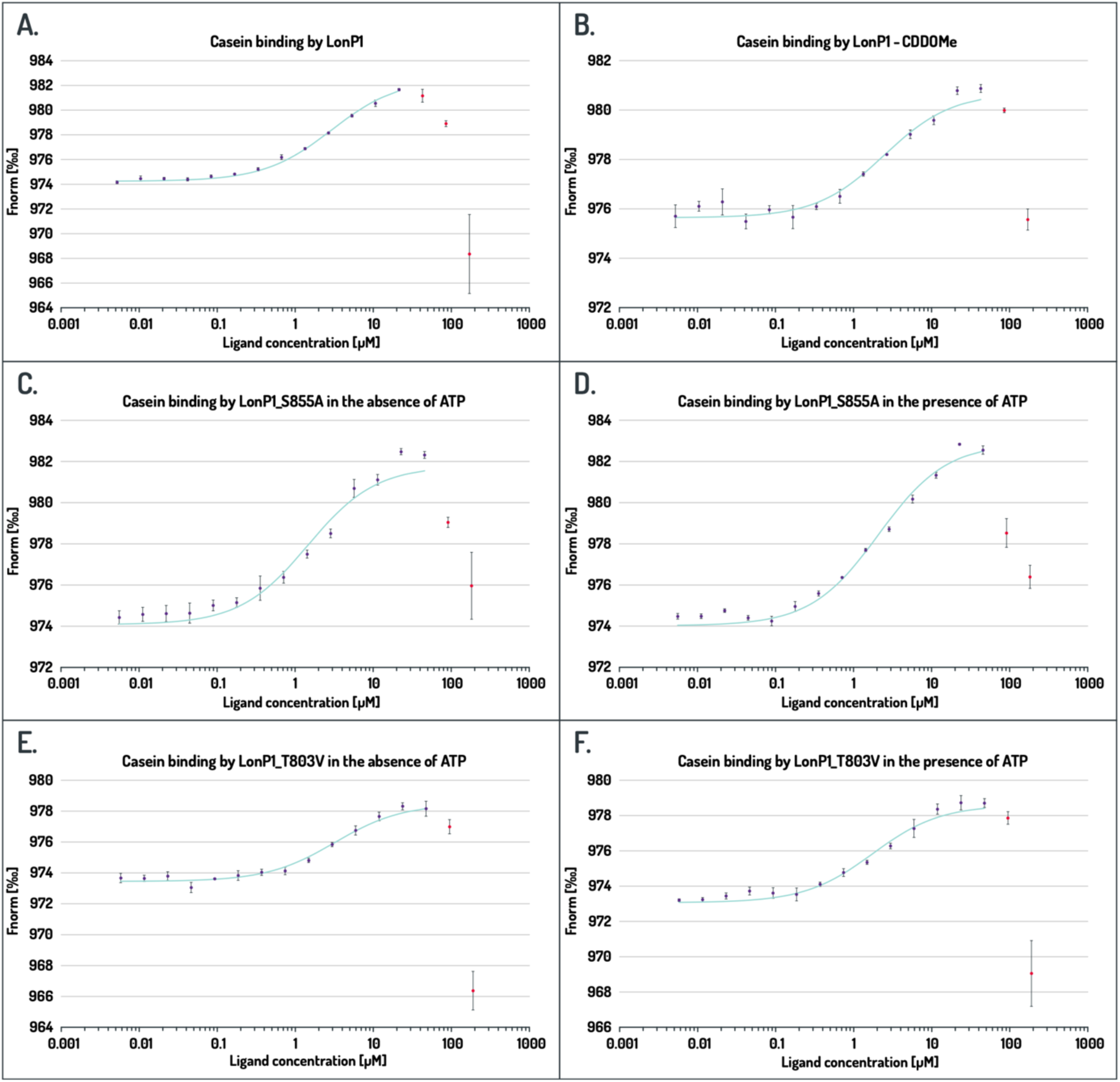
Substrate binding to LonP1 variants under different nucleotide and inhibition conditions. Binding of fluorophore-labelled LonP1 variants to the substrate protein casein was quantified by microscale thermophoresis. Changes in thermophoretic movement were recorded as fluorescence traces (Figure S5-S8), and the normalised fluorescence change (Fnorm) from the intial phase (0.5-1.5 s) was plotted against ligand concentration to obtain binding curves. Measurements were performed in triplicate and Fnorm was calculated for each trace (mean ± standard error). Masked out values are shown in pink. Apparent binding affinities were determined by non-linear regression (cyan). **A.** Casein binding to wild-type LonP1 in the absence of exogenous nucleotides: (KD = 2.95 ± 3.49 µM). **B.** Casein binding curve in the presence of the ATPase inhibitor CDDO-Me: (KD = 2.47 ± 1.4 µM). **C. and D.** Casein binding of the proteolytically inactive mutant LonP1_S855A in the absence (C: KD = 1.46 ± 3.81 µM) and presence (D: KD = 2.09 ± 1.79 µM) of ATP. **E. and F.** Casein binding of the protease mutant T803V in the absence (E: 3.20 ± 0.77 µM) and presence (F: 1.89 ± 0.94 µM) of ATP. **G. and H.** Casein titration to LonP1 inhibited with bortezomib in the absence (G: KD = 3.32 ± 1.95 µM) and presence (H: 2.07 ± 1.88 µM) of ATP.

**Figure S7:**
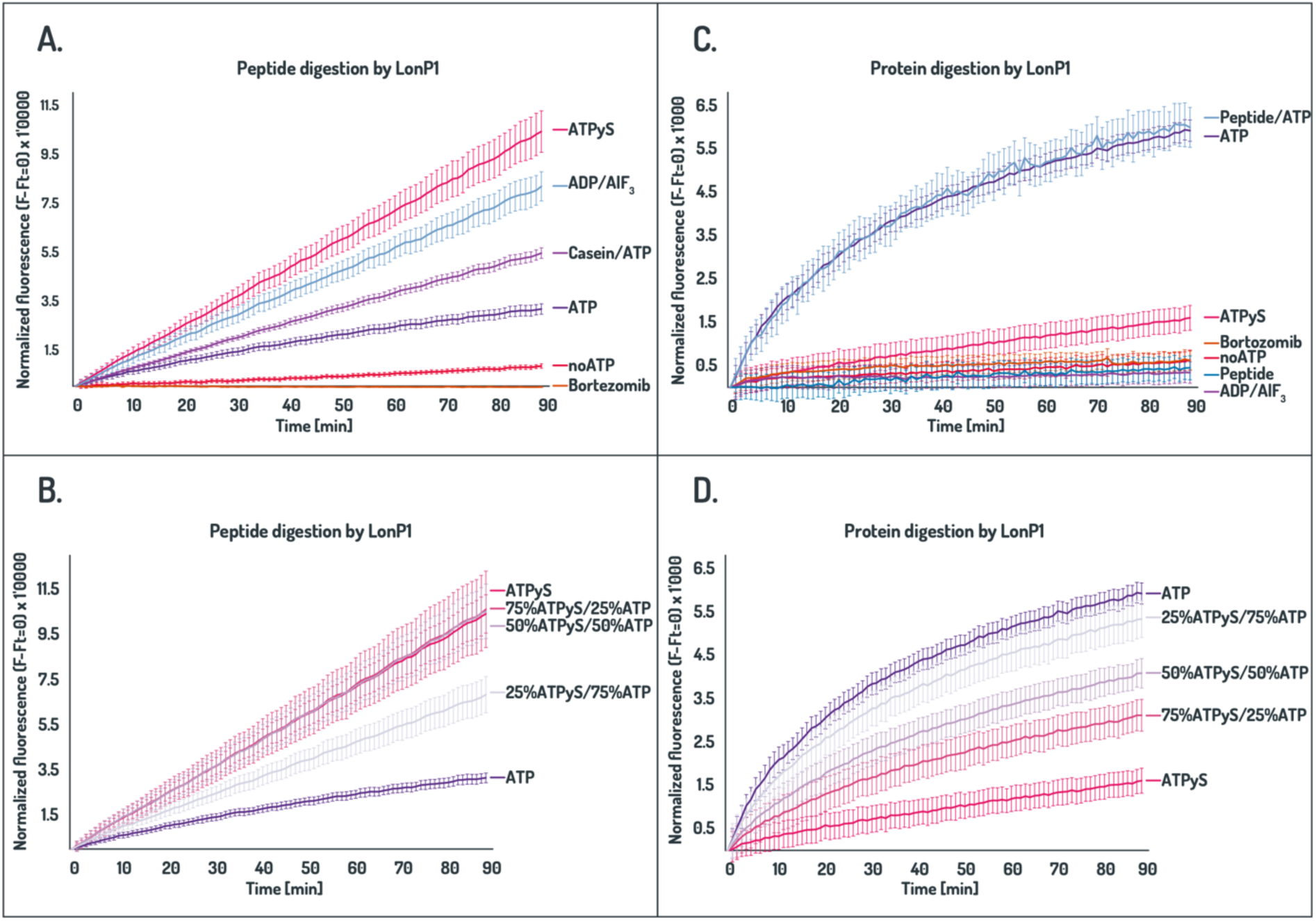
Digestion of a substrate peptide and protein by LonP1 under different nucleotide conditions. Proteolytic activity of LonP1 was evaluated using the fluorogenic peptide glutaryl-Ala-Ala-Phe-MNA and the protein substrate FITC-casein in the presence of ATP, ATPγS, and defined mixtures of both nucleotides. The peptide substrate is sufficiently small to diffuse into the catalytic cavity of LonP1, whereas FITC-casein requires active ATP-dependent unfolding and translocation prior to degradation. Substrate turnover was monitored as an increase in fluorescence over time (mean ± standard error). Initial degradation rates are indicated in grey. **A-B.** Time course of degradation of the fluorogenic peptide glutaryl-Ala-Ala-Phe-MNA under the indicated nucleotide conditions. **C-D.** Time course of degradation of the protein substrate FITC-casein under the indicated nucleotide conditions.

**Figure S8.**
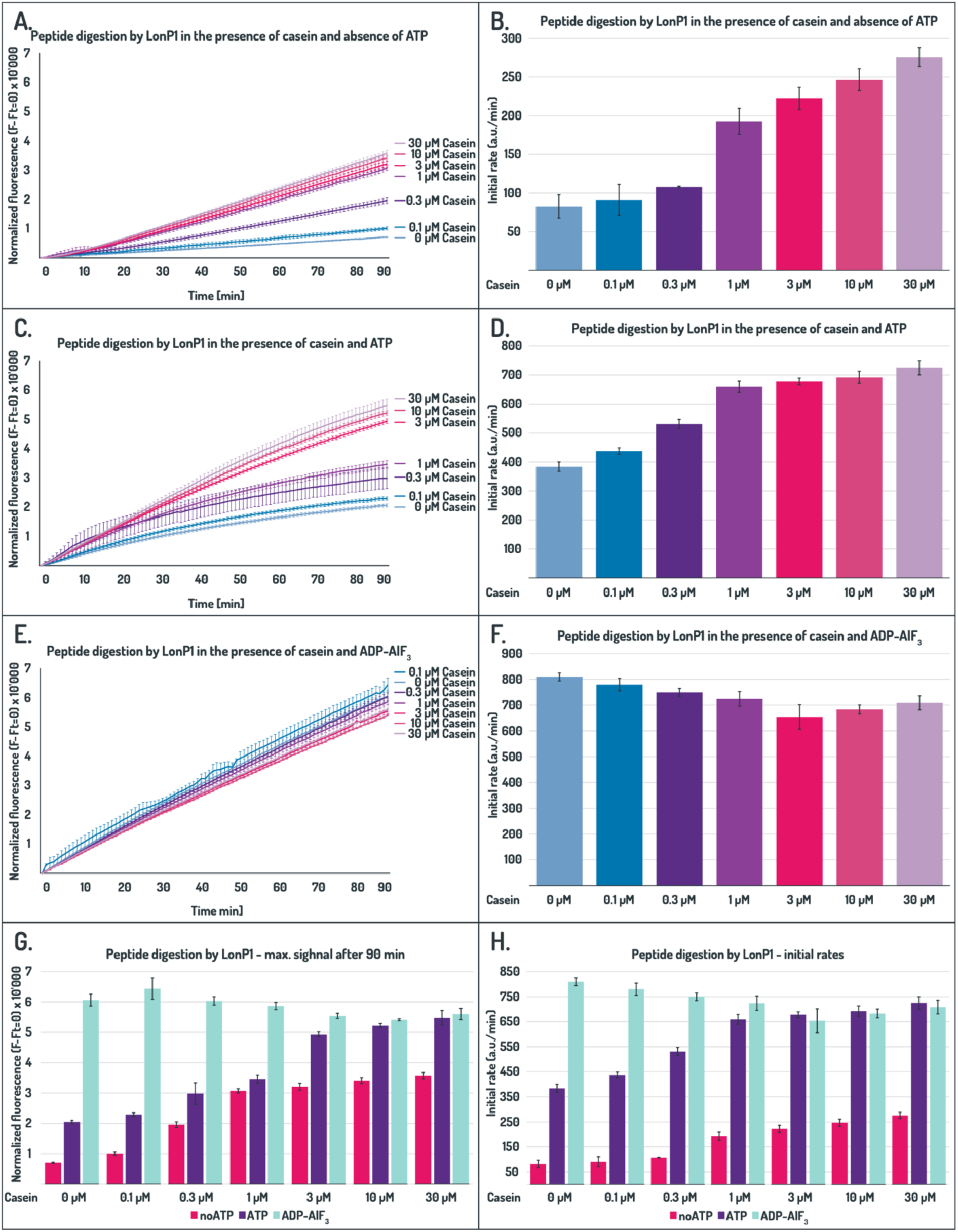
Cooperative modulation of LonP1 peptidase activity by protein substrate and nucleotides. Peptidase activity of LonP1 was analysed using a fluorogenic peptide, which can access the proteolytic chamber without mechanical unfolding. Casein concentration effects were analysed in the absence of nucleotides and in the presence of ATP or ADP·AlF₃. Substrate turnover was monitored as an increase in fluorescence over time at the respective casein concentrations (mean ± standard error) (A, C and E). To the right, the initial degradation rates determined from the linear phase of the respective fluorescence traces (4-8 min) are presented (mean ± standard error) (B, D and F). **A and B.** Peptide degradation in the absence of nucleotide. **C and D.** Peptide degradation in the presence of ATP (protease-competent state). **E and F.** Peptide degradation under ADP·AlF₃ conditions (transition-state mimic). **G.** Endpoint fluorescence after 90 min derived from the traces above. **H.** Combined initial degradation rates for comparison (B, D and F).

**Figure S9:**
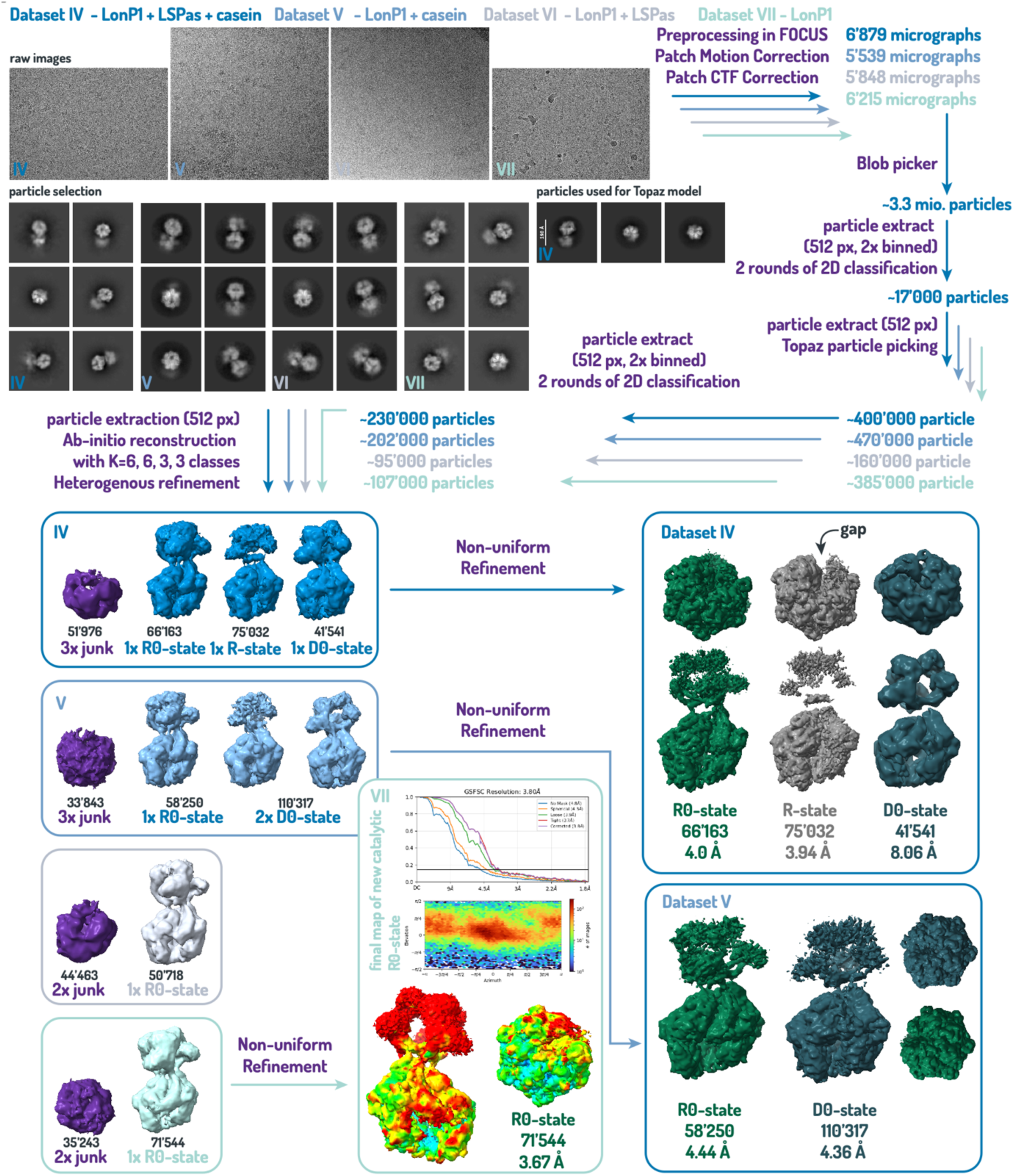
Data collection and 3D density calculation of LonP1 under different nucleotide-free conditions. Initial image processing, including motion correction, CTF estimation, particle picking, and 2D classification, was performed using closely matched parameters across all datasets to enable direct comparison. Differences between datasets became apparent during 3D classification. For dataset IV (LonP1 + LSPas + casein), three major particle classes were obtained that could be assigned to the canonical resting (R) state and two closely related conformations. All classes display a left-handed helical arrangement of the A-domains and one poorly ordered subdomain. However, the two newly observed conformations lack the inter-subunit gap characteristic of the open R-state (R0 and D1). Increasing the LonP1 concentration approximately ten-fold in dataset V (LonP1 + casein) shifted the particle distribution towards the closed D0-state, while the open R-state was no longer observed. In datasets lacking substrate protein (datasets VI and VII), only particles corresponding to the closed R0-state were detected. As dataset V still contained mitochondrial ssDNA (LSPas), whereas dataset VI did not, the emergence of the new conformations could be attributed to nucleotide-free conditions and substrate-dependent effects rather than to the presence of DNA. No additional density corresponding to LSPas could be detected.

**Figure S10:**
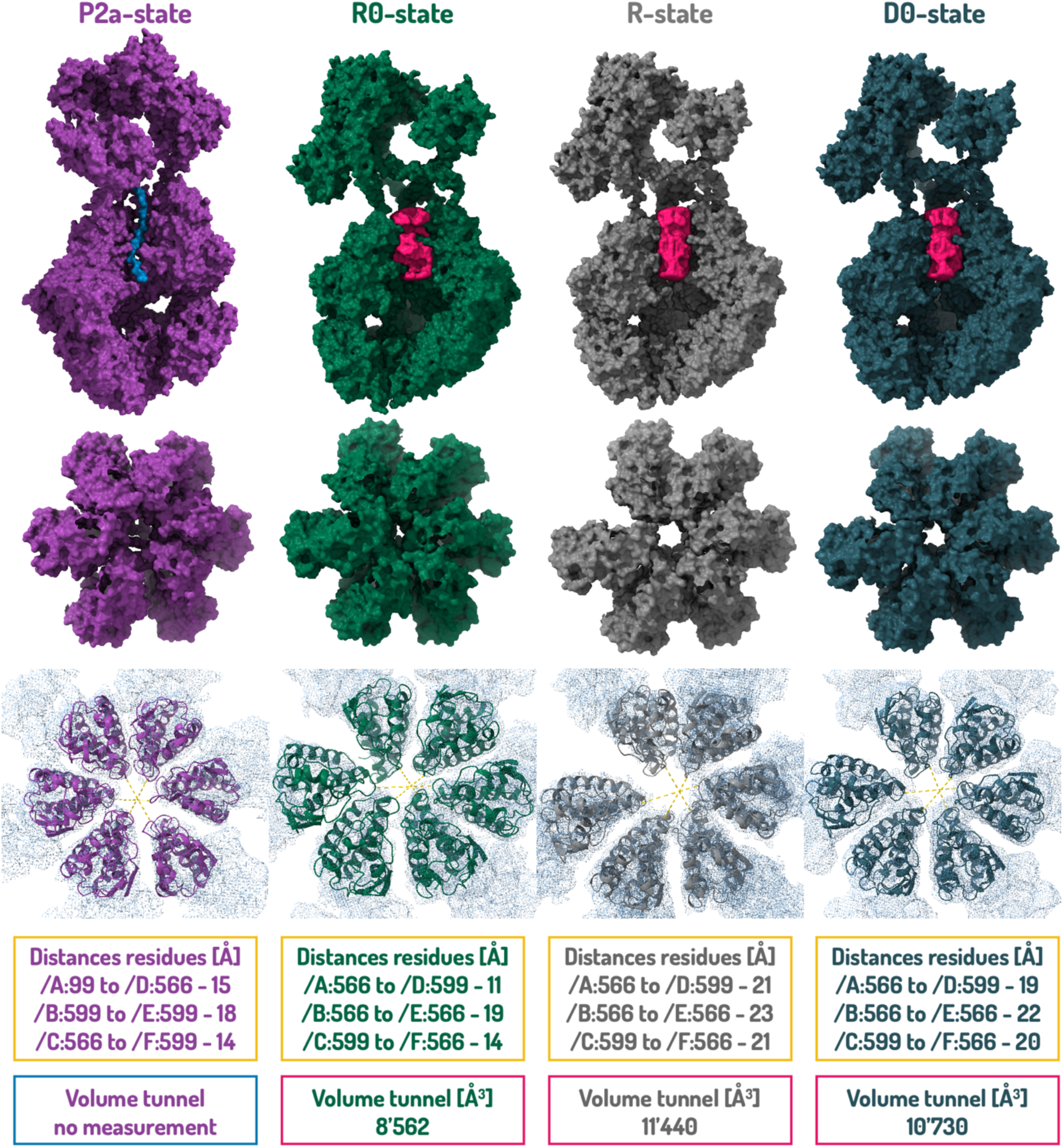
Quantitative analysis of the A-tunnel geometry in the different conformational states. Tunnel volumes were calculated using HOLLOW (Ho and Gruswitz, 2008), and distances between opposing loop regions lining the tunnel were measured in ChimeraX 1.10.1 (Goddard et al., 2018).

**Table S1:** Sample preparation Dataset I to IV.

| No. | Dataset | LonP1<br>[mg/ml,<br>μM] | LSPas<br>[μM] | Casein<br>[μM] | Quantifoil<br>Holey Carbon<br>Grids | Blotting conditions | Data acquisition |
| --- | --- | --- | --- | --- | --- | --- | --- |
| I | LonP1 +<br>LSPas +<br>casein | 0.9, 1.6 | 5.6 | 1.4 | 1.2/1.3 Cu200 | 3.5 μL reaction mix, 10 °C,<br>force 1, 2 sec, 100% relative<br>humidity | Dose: 54 e<sup>-</sup>/Å<sup>2</sup><br>Exposure: 4 s<br>Frames: 40 |
| II | LonP1 +<br>casein | 1, 1.7 |  | 10 | 2.1 Cu300 | 3.5 μL reaction mix, 10 °C,<br>force 1, 3 sec, 100% relative<br>humidity | Dose: 50 e<sup>-</sup>/Å<sup>2</sup><br>Exposure: 8.4 s<br>Frames: 40 |
| III | LonP1 +<br>LSPas | 0.8, 1.4 | 5 |  | 2.1 Cu300 | 3.5 μL reaction mix, 10 °C,<br>force 1, 3 sec, 100% relative<br>humidity | Dose: 50.3 e<sup>-</sup>/Å<sup>2</sup><br>Exposure: 4.3 s<br>Frames: 40 |
| IV | LonP1 | 0.7, 1.2 |  |  | 1.2/1.3 Cu200 | 3.5 μL reaction mix, 10 °C,<br>force 1, 3 sec, 100% relative<br>humidity | Dose: 40 e<sup>-</sup>/Å<sup>2</sup><br>Exposure: 2.2 s<br>Frames: 40 |

**Table S2:**
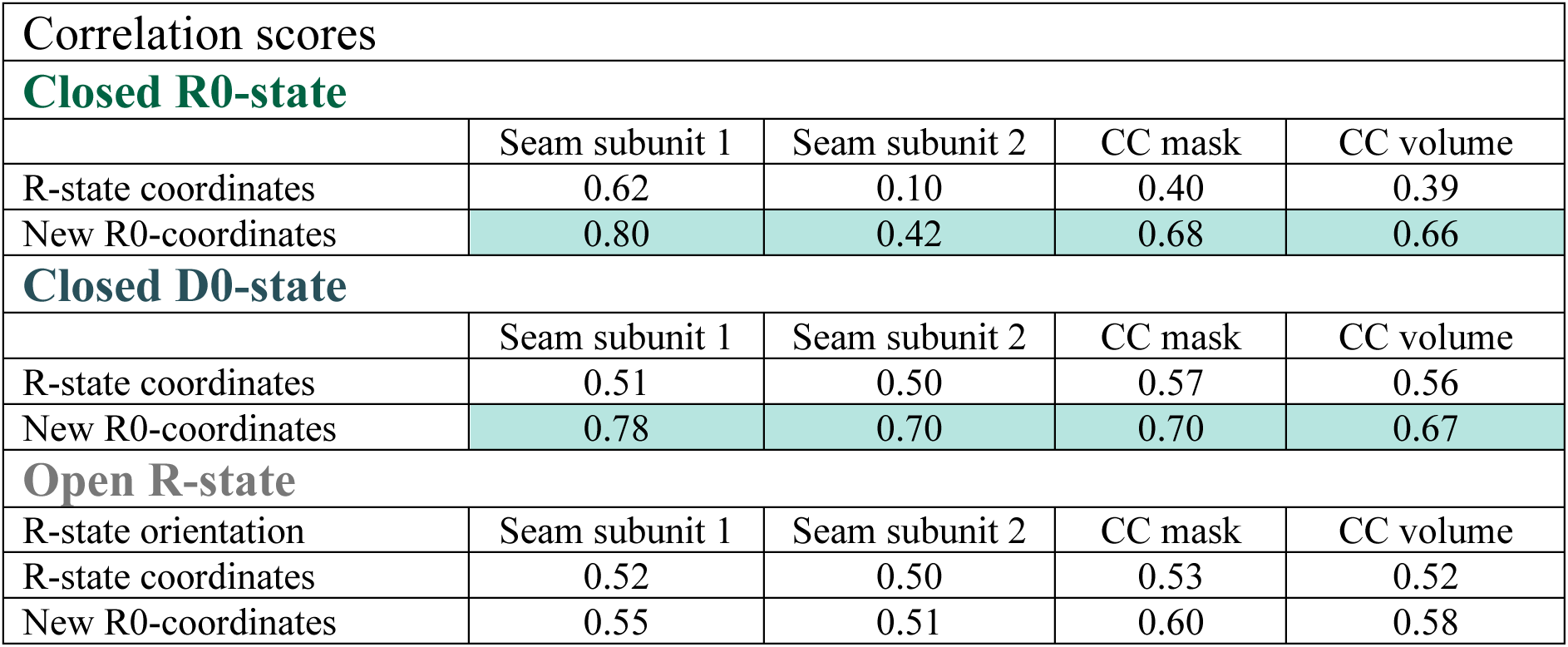
Model-map correlation analysis distinguishes open and closed resting conformations of LonP1. To discriminate between the three observed resting conformations of LonP1 (open R, closed R0, and closed D0), cryo-EM maps were aligned using the prominently unstructured subunit as a reference landmark, enabling consistent analysis of the adjacent seam region. An initial atomic model was generated for the more distinct closed R0-state. This model, together with available coordinates of the canonical open R-state (PDB: 7NGL), was subsequently used for comparative fitting. For each map, the A- and P-domains of four subunits located opposite the seam were placed into the density and subjected to rigid-body refinement. The N-terminal domains were excluded from refinement due to their pronounced rotational variability between states. Following refinement, the complete A/P-domain assemblies were superimposed onto the refined coordinates, and model-map correlation coefficients (CC) were determined specifically for the seam subunits. This analysis enabled validation of structural differences between the closed and open resting conformations (cyan). While the open R-state is characterised by a pronounced inter-subunit gap at the seam, the local resolution in this region is limited, resulting in comparatively small differences in correlation scores between the fitted models.

**Table S3:** Refinement statistics.

| Parameter | R0-state | D0-state |
| --- | --- | --- |
| **MODEL COMPOSITION & GEOMETRY** |  |  |
| Chains | 6 | 12 |
| Atoms (with Hydrogens) | 74'807 (37'787) | 74'807 (37'787) |
| Residues (Protein / Nucleotide) | 4'656 / 0 | 4'656 / 0 |
| Water molecules | 0 | 0 |
| Ligands | ADP (6) | ADP (6) |
| **Bonds (RMSD)** |  |  |
| Length (Å) (# > 4σ) | 0.004 (0) | 0.006 (0) |
| Angles (°) (# > 4σ) | 0.648 (19) | 0.950 (11) |
| **Validation Scores** |  |  |
| MolProbity score | 2.01 | 1.32 |
| Clash score | 9.41 | 2.10 |
| Rotamer outliers (%) | 1.56 | 0.00 |
| Cβ outliers (%) | NA | NA |
| CaBLAM outliers (%) | 3.12 | 2.56 |
| **Ramachandran Plot (%)** |  |  |
| Favored | 94.60 | 95.06 |
| Allowed | 5.31 | 4.94 |
| Outliers | 0.09 | 0.00 |
| **Rama-Z (Plot Z-score, RMSD)** |  |  |
| Whole | -1.17 (0.12) | -1.66 (0.11) |
| Helix | 0.40 (0.11) | -0.22 (0.11) |
| Sheet | -2.36 (0.21) | -2.54 (0.19) |
| Loop | -1.48 (0.14) | -1.45 (0.13) |
| **Peptide Plane (%)** |  |  |
| Cis proline / general | 0.0 / 0.0 | 0.0 / 0.0 |
| Twisted proline / general | 0.0 / 0.0 | 0.0 / 0.0 |
| **B-factors / Atomic Displacement Parameters (Å²)** |  |  |
| Iso/Aniso (#) | 37'020 / 0 | 37'020 / 0 |
| Protein (min / max / mean) | 123.39 / 996.90 / 411.08 | 161.87 / 1031.34 / 544.19 |
| Ligand (min / max / mean) | 132.20 / 640.77 / 271.07 | 182.03 / 425.04 / 246.59 |
| **Occupancy** |  |  |
| Mean | 1.00 | 1.00 |
| occ = 1 (%) | 99.98 | 99.98 |
| **DATA STATISTICS & RESOLUTION** |  |  |
| Box Lengths (Å) | 146.63, 142.24, 220.38 | 145.96, 138.58, 218.94 |
| Box Angles (°) | 90.00, 90.00, 90.00 | 90.00, 90.00, 90.00 |
| Supplied Resolution (Å) | 3.8 | 4.2 |
| **Resolution Estimates (Å) [Masked / Unmasked]** |  |  |
| d FSC (half maps; 0.143) | --- / --- | --- / --- |
| d 99 (full/half1/half2) | 3.0 / 2.9 | 3.2 / 3.1 |
| d model | 2.9 / 2.9 | 3.1 / 3.2 |
| d FSC model (0/0.143/0.5) [Masked] | 3.6 / 3.8 / 4.5 | 3.9 / 4.3 / 7.0 |
| d FSC model (0/0.143/0.5) [Unmasked] | 3.6 / 4.1 / 4.8 | 4.2 / 4.5 / 7.8 |
| Map min / max / mean | -0.13 / 0.34 / 0.01 | -0.05 / 0.22 / 0.01 |
| **MODEL VS. DATA CORRELATION (CC)** |  |  |
| CC (mask) | 0.77 | 0.80 |
| CC (box) | 0.87 | 0.90 |
| CC (peaks) | 0.65 | 0.69 |
| CC (volume) | 0.76 | 0.79 |
| Mean CC for ligands | 0.80 | 0.88 |

